# Auxiliary ATP binding sites power rapid unwinding by RecBCD

**DOI:** 10.1101/210823

**Authors:** Rani Zananiri, Vera Gaydar, Dan Yahalom, Omri Malik, Sergei Rudnizky, Ariel Kaplan, Arnon Henn

## Abstract

RecBCD, responsible for the initiation of double stranded break repair in bacteria, is a processive DNA helicase with an unwinding rate approaching ∼1,600 bp·s^−1^. The mechanism enabling RecBCD to achieve such fast unwinding rate is not known. We employed a combination of equilibrium and time–resolved binding experiments, and ensemble and single molecule activity assays to uncover the molecular mechanism underlying RecBCD’s rapid catalysis. We report the existence of auxiliary binding sites, where ATP binds with lower affinity and with distinct chemical interactions as compared to the known catalytic sites. The catalytic rate of RecBCD is reduced both by preventing and by strengthening ATP binding to these sites, suggesting that the dynamics of ATP at these sites modulates the enzyme’s rate. We propose a model by which RecBCD achieves its fast unwinding rate by utilizing the weaker binding sites to increase the flux of ATP to its catalytic sites.

## INTRODUCTION

Double Strand Breaks (DSBs) are the severest type of DNA damage in all kingdoms of life. Ubiquitous repair mechanisms for DSBs are found in every living organism, wherein helicases play an essential role. In prokaryotes, members of the RecBCD family initiate unwinding of the DNA at the DSB site in preparation for strand invasion, which is essential for the homologous recombination DSB repair pathway^1^. RecBCD is a heterotrimer consisting of one copy each of the RecB and RecD DNA translocases and helicases of opposing unwinding polarity^2, 3^. The RecC subunit “staples” the RecB and RecD subunits^4^, harbors the pin domain that splits the duplex and sends each strand down separate channels^4^, and recognizes the Chi sequence^5, 6^.

Previous studies have provided a wealth of knowledge on RecBCD’s biochemistry and structure^1, 7, 8, 9, 10, 11^ and revealed important features of the unwinding process, including its initiation, unwinding kinetic step size, and inter-subunit regulation^11, 12, 13, 14, 15^. Collectively, these studies have produced a working model for its mechanism of coupled unwinding and translocation. The opposing translocation polarities of RecB and RecD synchronize the translocation of the RecBCD complex along the duplex DNA^3, 7^. However, the unwinding rates of RecB and RecD are different, with RecD translocating faster than RecB under physiological conditions^7, 11, 16^, an asymmetry that results in a single-stranded loop that accumulates on the 3’-ended strand^7, 11^. Recognition of the Chi sequence during processive unwinding causes a major conformational change in RecBCD^16, 17^, which modulates the nuclease activity of the C-terminus domain of RecB^18^, switching it from endonucleolytically nicking primarily the 3’-ended strand to nicking primarily the 5’-ended strand^19, 20^. Finally, RecBCD promotes recruitment of the DNA strand-exchange protein RecA onto the 3’-end of the ssDNA tail^21^.

RecBCD is a highly processive helicase exhibiting an exceptionally high unwinding rate of ∼ 1,600 base pairs (bp) per second (s^−1^) ^22^. Given that it consumes two ATP molecules per DNA bp unwound^23, 24^, this amasses to a lower limit of 3,200 hydrolyzed ATPs s^−1^ RecBCD^−1^, meaning that RecBCD is able to complete 3 ATPase cycles in less than a millisecond. Thus, while previous studies focused mainly on delineating the order of events that encompass the unwinding and translocation reaction, we believe that it is also essential to elucidate what are the specific properties allowing RecBCD to achieve its rapid catalytic cycle.

In this work, we characterized the binding of nucleotides to RecBCD and RecBCD× DNA complexes, and the unwinding and translocation activity of RecBCD on different DNA substrates. Our results reveal the existence of auxiliary, lower affinity ATP binding sites in RecBCD, in addition to the known catalytic binding sites. These sites, likely located in RecC, are distinct in their affinity and kinetics of binding, and in their sensitivity to ATP-analogues and salt. Using a real-time, single-turnover, anisotropy-based DNA unwinding assay and a novel single-molecule, optical tweezers assay, we find that the lower affinity sites contribute significantly to both the ATPase rate and the unwinding velocity, but only at ATP concentrations that are much higher than those required to fully saturate the canonical binding sites in both subunits. In addition, both binding and unbinding of ATP at these sites is required to achieve the full catalytic rate. A model where ATP binding to the auxiliary binding sites serves to increase the flux of ATP to the catalytic sites fully recapitulates our binding and activity measurements.

## RESULTS

### RecBCD possesses strong and weak nucleotide binding sites

The ADP and ATP affinities of RecBCD were determined using fluorescent nucleotide analogues, mantADP and mantAMPpNp, respectively. The binding is measured as a Förster Resonance Energy Transfer (FRET) signal between the intrinsic tryptophans (λ_ext_ = 280 nm) and the mant-nucleotides (λ_em_ = 436 nm). Remarkably, the pattern by which RecBCD binds to mant-nucleotides follows a biphasic behavior as a function of the nucleotide concentration (Figs. 1A & 1B). An alternative characterization of ADP binding by competition of unmodified ADP to a pre-equilibrated RecBCD·mantADP complex, yielded a similar biphasic pattern (Supplementary Fig. 1). At first sight, one might attribute each of the binding phases to a single nucleotide binding site in each one of RecBCD’s catalytic sites, one in RecB and one in RecD. However, the equations describing binding of a ligand to a macromolecule containing two binding sites cannot result in a biphasic pattern of this kind. The reader is referred to the Supplementary Information (Supplementary Results and Supplementary Fig. 2) for a comprehensive analysis regarding the analysis of the binding curve isotherms. Briefly, the biphasic curve can be decomposed into two phases: one hyperbolic with strong affinity, and one sigmoidal with weak affinity. This decomposition emphasizes why two sites are insufficient to give rise to a biphasic pattern: The hyperbolic phase can be the result of binding to a single site or multiple non-cooperative sites. However, the sigmoidal phase can only be observed if there are at least two cooperative sites that are distinct from the site/s of the first phase. This accumulates to at least three nucleotide binding sites within RecBCD. Hence, as a phenomenological characterization, we describe the binding isotherms as the sum of two Hill equations (Eq. 1, Methods). The first Hill curve describes stronger affinity, but weakly cooperative binding and is characterized with a macroscopic equilibrium constant *K*_s_ and a Hill coefficient *n*_s._ The second Hill curve describes weaker affinity, but cooperative, nucleotide binding sites, with an overall macroscopic equilibrium constant *K*_w_ and Hill coefficient *n*_w_. The results of fitting such a model to the binding of mant-nucleotides are summarized in Supplementary Table 1.

**Figure 1:**
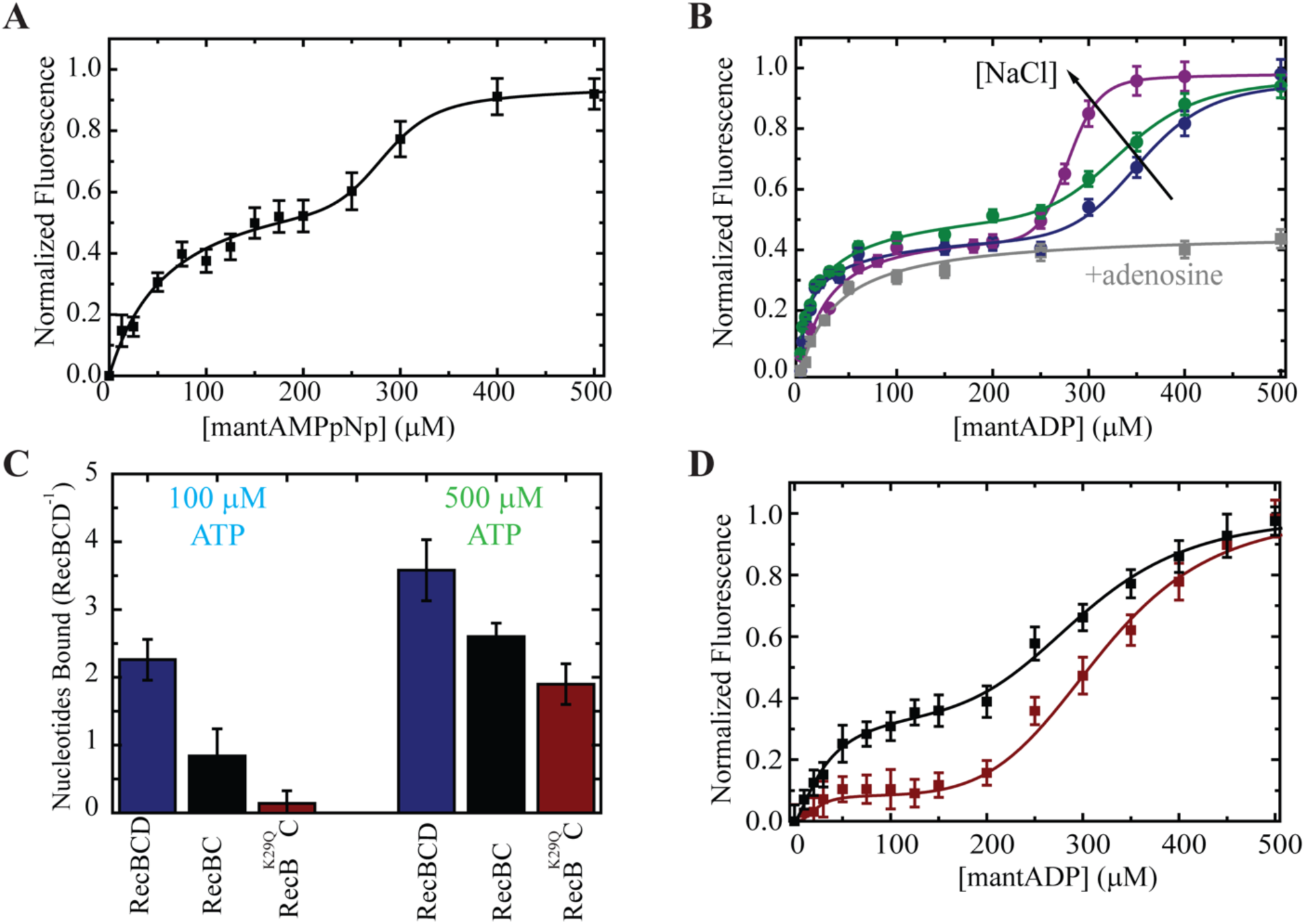
Equilibrium nucleotide binding to RecBCD. **A.** Titration curves of mantAMPpNp to RecBCD exhibit a biphasic pattern reaching saturation at ∼500 μM nucleotide concentration. Lines show the best fit to Eq. 1 (Methods). Data shown as mean ± s.e.m., n = 3. **B.** Salt and adenosine dependence of mantADP binding to RecBCD. Blue, green and purple correspond to 75, 200 and 300 mM NaCl, respectively, and gray represents the binding curve in the presence of 75 mM NaCl and 2 mM adenosine. Data shown as mean ± s.e.m., n = 3. Lines are best fits to Eq. 1 (Methods). **C.** Number of ADP molecules bound to RecBCD (blue), RecBC (black) and RecB^K29Q^CD (red), measured using equilibrium dialysis, for low (∼100 µM, left) and high (∼500 µM, right) nucleotide concentrations. Data shown as mean ± s.e.m.; n = 2 and 4, for low and high concentrations, respectively. **D.** Titration curves of mantADP to RecBC (black) and RecB^K29Q^CD (red). Lines show the best fit to Eq. 1 (Methods). Data shown as mean ± s.e.m., n = 3.

**Figure 2:**
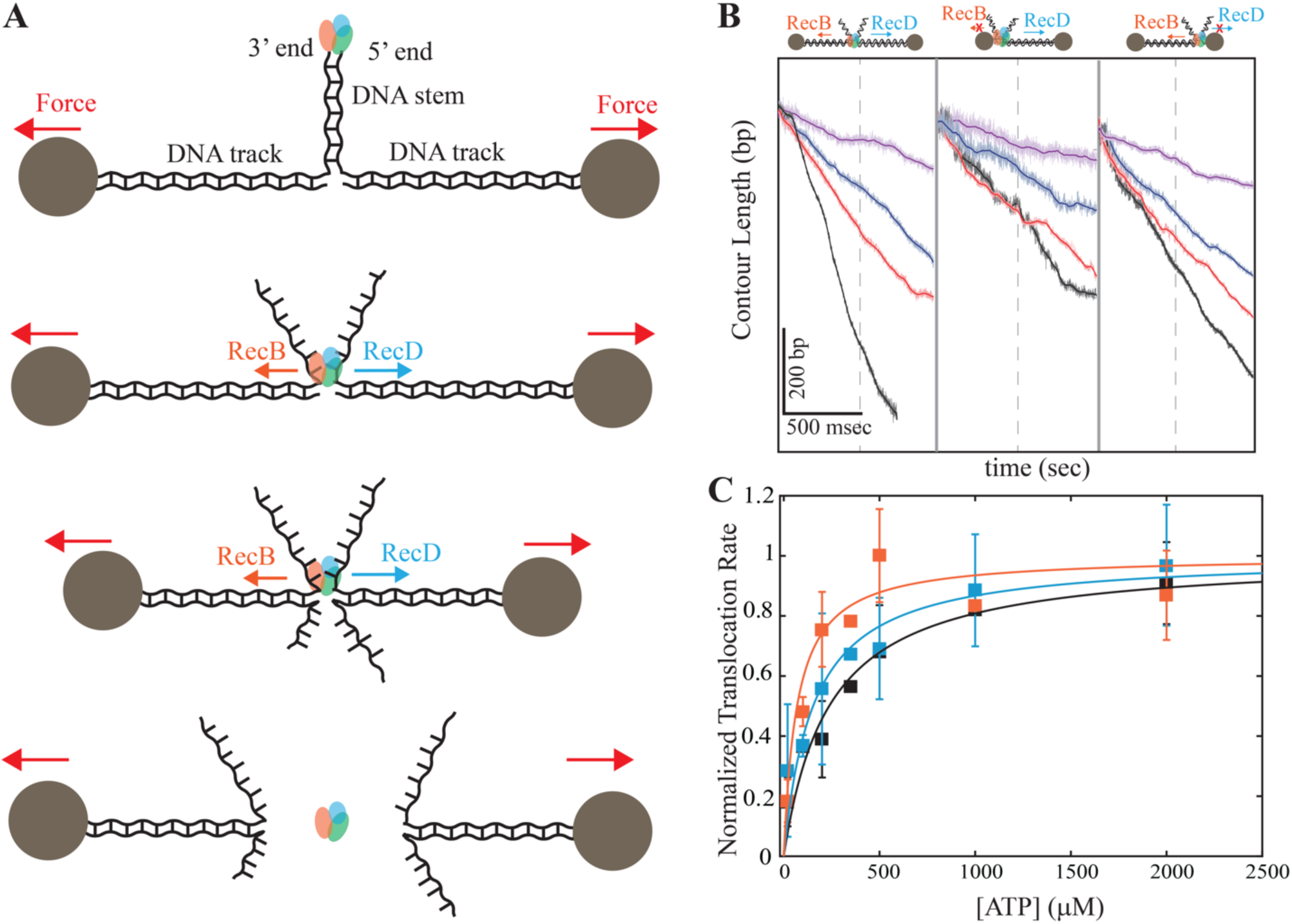
Single molecule measurements of translocation by RecBCD and its individual subunits. **A**. Schematic representation of the experimental optical tweezers setup. RecBCD binds to and translocates on a DNA stem connected to beads through DNA ‘tracks’. Upon reaching the fork, the helicase subunits translocate in different directions due to their opposing polarities, shortening the tether length. The force increases up to a point where RecBCD dissociates from the construct. **B**. Representative traces of translocations on three constructs probing both translocases (left, symmetric tracks of 600 bp), RecD (middle, asymmetric tracks of ∼30 nt/∼4000 bp), and RecB (right, asymmetric tracks of ∼4000 bp/∼30 nt) at different ATP concentrations (black, 2 mM; red, 500 µM; blue 200 µM; purple, 50 µM). Light colors display the raw, 2.5kHz data, and bold lines a moving average of 100 points. **C**. Translocation rates versus [ATP] for both translocases (black), RecD (blue) and RecB (orange) in the force range 10-15 pN. Solid lines through the data points are fits to hyperbolic curves. Data shown as mean ± s.e.m., normalized to the maximal rate according to the fit. The number of traces used in the analysis is summarized in Supplementary Table 2).

### RecBCD binds at least four nucleotides

The equilibrium binding experiments reveal the existence of additional binding sites, but not their *n*-value (number of sites). Moreover, they do not reveal the relative stoichiometry between sites corresponding to the different binding phases, since these sites may involve a different binding mode and hence a different FRET efficiency. Hence, to quantify the number of nucleotide binding sites on RecBCD we employed equilibrium dialysis, a first-principle and model-free method to study binding of a ligand to a macromolecule. In this assay, a semipermeable membrane separates the ligand, i.e. ADP or AMPpNp, from RecBCD (apo or pre-bound to DNA, Supplementary Table 7, 8) and the samples are allowed sufficient time to equilibrate. The concentration of ligands in the RecBCD-free compartment, before and after equilibration, is then used to calculate the number of ligands bound to RecBCD. The experiments were performed at two nucleotide concentrations: a low concentration (200 µM pre-equilibration; Supplementary Fig. 3A), where we aimed to saturate only the hyperbolic binding phase in our measured isotherms, and a high concentration (1 mM nucleotides pre-equilibration; Supplementary Fig. 3B), at which we aimed to saturate both binding phases. In the low nucleotide concentration regime, RecBCD binds *n* = 2 ATP molecules/RecBCD (RecBCD·ADP: *n* = 2.26 ± 0.30; RecBCD·AMPpNp: *n* = 1.94 ± 0.15; average of two independent measurements ± s.e.m.; Figure 1C and Supplementary Fig. 3D). However, in the high nucleotide concertation regime we found *n* = 3.5-4 ATP molecules/RecBCD for all the RecBCD complexes we measured (RecBCD·ADP: *n* = 3.58 ± 0.45; RecBCD·AMPpNp: *n* = 3.48 ± 0.70; RecBCD·ohDNA·ADP: *n* = 3.99 ± 0.50; average of four independent measurements ± s.e.m.). We note that the high concentration experiments were performed at sub-stoichiometric concentrations (limited by [RecBCD]) and may not represent full saturation of all ligand binding sites. Hence, our results provide a lower limit on the number of binding sites, and suggest that the strong binding phase accounts for two binding sites and the weak binding phase represents binding to at least two additional sites.

**Figure 3:**
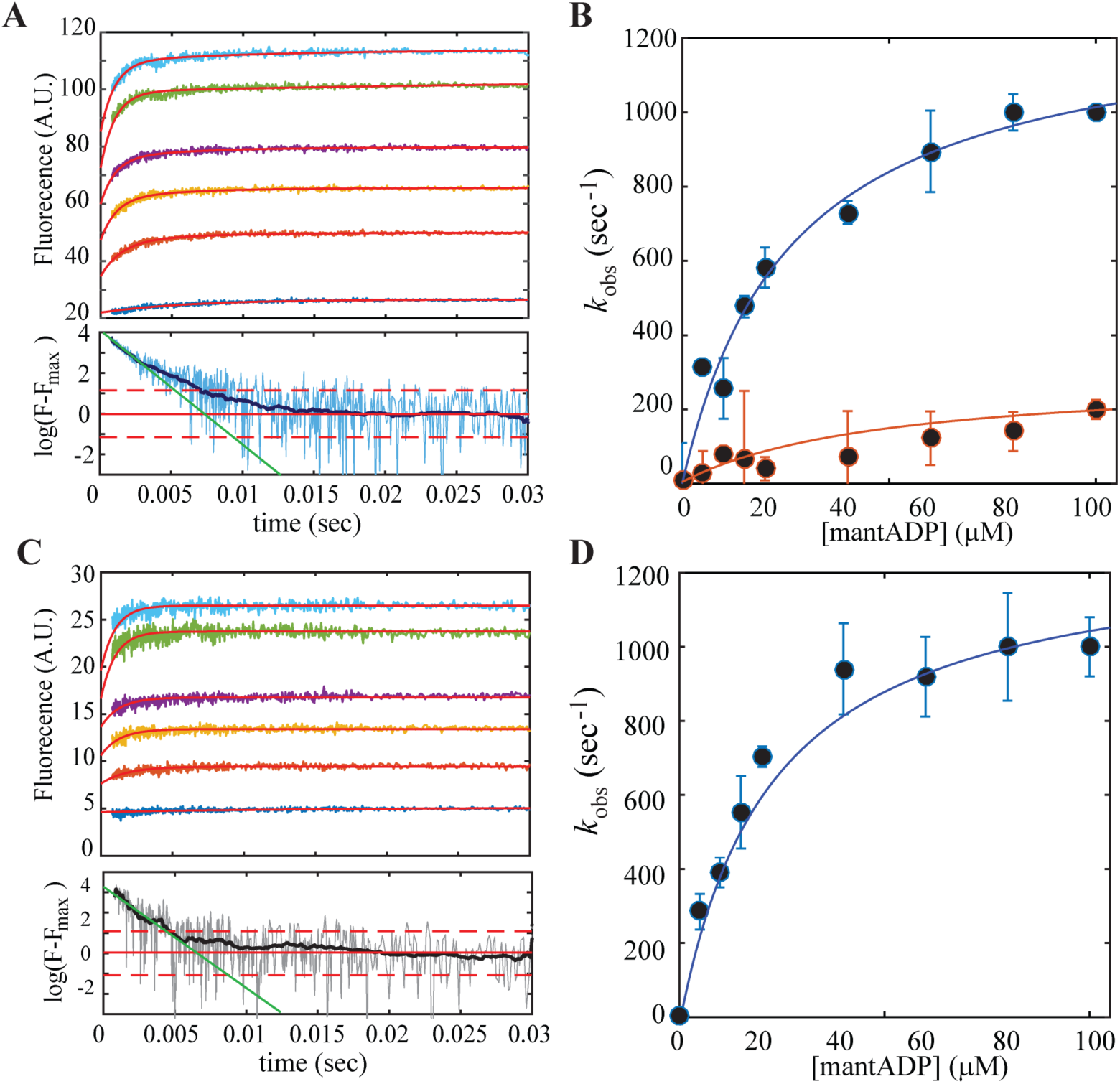
Transient kinetics of mantADP binding to RecBCD. **A. Top:** Time courses of mantADP binding upon rapid mixing of RecBCD (2 µM, post-mixing) with mantADP (0-100 µM, lower to upper, respectively). The red lines through the data are the best global fit to a double exponential function (Methods). **Bottom:** Log-scale representation of the normalized intensity highlights the deviation from a single exponential. The light-blue line shows raw data of 60 µM and the dark-blue filtered data (Methods). Red horizontal lines indicate mean (full line) and standard deviation (two dashed lines) of the noise. The green line is the best linear fit for the first 4 msec. In the time window from time t=0 to the time when the signal equals the noise, up to a standard deviation, the data shows deviation from a single exponential decay. **B.** Dependence of 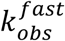 (Blue) and 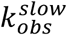 (Orange) on [mantADP], both displaying hyperbolic dependencies. Data shown as mean ± s.e.m., n = 6-7. The solid line through the data points are best hyperbolic fits. **C. Top:** Time courses of mantADP binding upon rapid mixing of RecBCD (2 µM, post-mixing) with mantADP (0-100 µM, lower to upper, respectively) in the presence of 2mM adenosine. The red lines through the data are the best global fit to a single exponential function (Methods). **Bottom:** Log-scale representation of the normalized intensity highlights a consistency with a single exponential. The gray line shows raw data of 60 µM and the black filtered data (Methods). Red horizontal lines indicate mean (full line) and standard deviation (two dashed lines) of the noise. The green line is the best linear fit for the first 4 msec. **D.** Dependence of *k*_*obs*_ on [mantADP] in the presence of adenosine, displaying a hyperbolic dependence. Data shown as mean ± s.e.m., n = 6-7. The solid line through the data points are best hyperbolic fits.

### Weak binding sites are likely located at the RecC subunit

The biphasic binding curves and the equilibrium dialysis experiments support the existence of additional binding sites for ATP, other than the canonical catalytic ones. Hence, to elucidate the potential location of such binding sites, we computationally scanned the protein surface (subunits B, C and D of RecBCD) for putative binding sites utilizing small molecules as probes according to the FTMap method^25^. We found that, beyond the catalytic sites in RecB and RecD, several such clusters were found, in particular large clusters located in RecC (Supplementary Fig. 4). Motivated by these findings, we assayed mant-ADP binding to RecBC (i.e. a complex lacking the D subunit). Fig. 1D shows that RecBC binds to mant-nucleotides in a biphasic manner as well. Interestingly, since the affinity of the strong binding phase decreased in the RecBC case, with only one catalytic site, as compared with RecBCD (*K*_s_ increased from ∼15 µM to ∼23 µM) while the weak binding phase remained similar, these results suggest that the strong binding phase corresponds to binding to the catalytic sites, and the weak binding phase to the additional sites, which are likely located in the RecC subunit. Remarkably, when we repeated these experiments with the catalytically-deficient mutant RecB^K29Q^D, we found that the binding isotherms show a significant reduction in the strong binding phase (Fig. 1D). In addition, equilibrium dialysis experiments showed practically no binding in the low nucleotide concentration regime and similar binding as for RecBC in the high nucleotide concentration regime (Fig. 1C), lending further support for our assignment of the binding sites.

**Figure 4:**
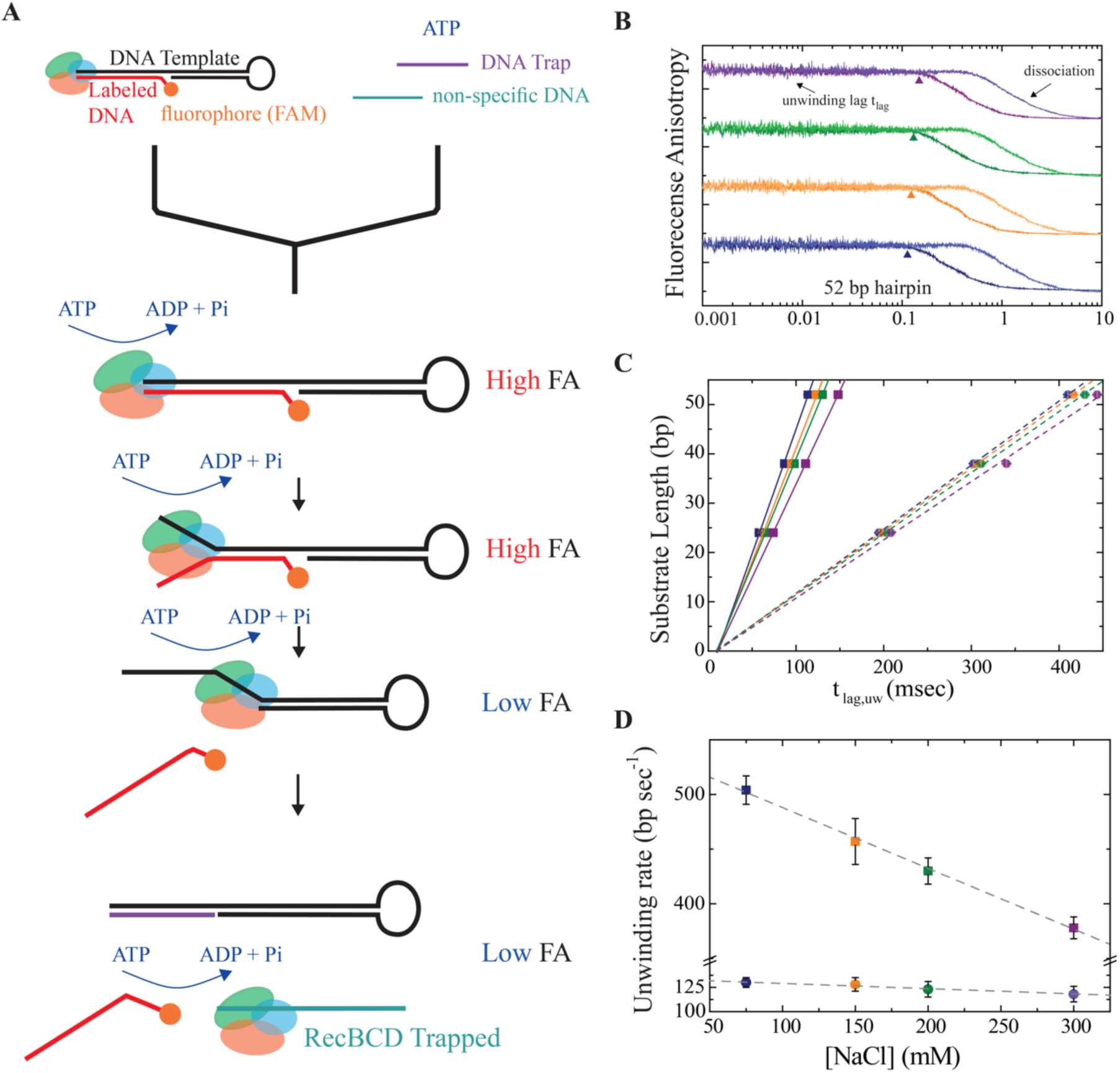
Rapid mixing, fluorescence anisotropy measurements of real time unwinding by RecBCD. **A.** Schematic description of the experimental setup. A pre-bound RecBCD/labeled-DNA complex is mixed with ATP and DNA traps designed to capture the released RecBCD and DNA template. A high FA phase corresponds to the unwinding phase, while a lower FA corresponds to the dissociation of RecBCD from the DNA template. **B.** FA time courses of unwinding reactions of RecBCD× hpDNA (52bp; 250 nM, post-mixed) with ATP (250 μM, strong colors and 100 μM, light colors). Time courses were vertically shifted for clarity. Difference colors indicate time traces at 75mM (blue), 150 mM (orange), 200 mM (green), and 300 mM (purple) NaCl concentrations. **C.** The unwinding lag duration as a function of the substrate length at high (squares) and low (circles) ATP concentrations. [NaCl] color coding as in B. Data shown as mean ± s.e.m., n = 11-14 for each point. Lines through the data are best linear fits. **D.** Unwinding rates as a function of [NaCl], as calculated from the slopes in Figure 4C, for high (squares) and low (circles) [ATP]. Data shown as mean ± s.e.

### ATP binds the strong and weak sites through distinct chemical interactions

To investigate whether the biochemical properties of the additional nucleotide binding sites are similar to the canonical ones, we measured the binding isotherms as a function of [NaCl]. Fig. 1B shows mantADP binding to RecBCD as a function of increasing [NaCl], and reveals that salt affects mainly the second phase of the binding curve, with an undistinguishable effect on the first binding phase (results of the fitting are summarized in Supplementary Table 1). This suggests that the strong ATP binding sites are largely not influenced by electrostatic interactions. However, the occupancy of the weak sites increases with higher [NaCl], indicating that binding at these sites is largely mediated by hydrophobic interactions, which are strengthened by salt^26^. We then examined the effect of adenosine nucleosides (nucleotides without the phosphate group) on mantADP binding to RecBCD. Excitingly, adding 2 mM adenosine specifically inhibited mantADP binding to the weak binding sites, resulting in a hyperbolic binding curve with similar binding affinity to the strong sites, *K* = 46 ± 6 μM (Fig. 1B), as the one measured in the first phase in the absence of adenosine. Taken together, these results indicate that the chemical nature of the strong and weak binding sites is different, with ATP interacting with the weak binding sites mainly through the base and sugar moiety of the nucleotide. Furthermore, they suggest that adenosine and salt can be used as experimental tools to specifically modulate one set of binding sites in RecBCD.

### RecBCD’s individual subunits are active at similar ATP concentrations

If RecB and RecD have significantly different affinities towards nucleotides, one would expect to find that, at an intermediate [ATP], one subunit will be fully active while the second will not. Hence, to correlate between our nucleotides binding model, of strong and weak binding sites, to the catalytic activity of RecBCD’s subunits, it is important to study the activity of RecBCD’s individual subunits. Importantly, since the biphasic binding curves could arise from cooperativity among the subunits, which can be diminished if mutations are introduced, we designed an experimental setup aimed to study the activity of the subunits in the context of a wildtype (WT) RecBCD, using a novel single-molecule optical-tweezers assay. Fig. 2A shows a schematic representation of the experiment: The DNA construct consists of a stem with a blunt end attached to two dsDNA “tracks”, each containing a specific tag at its 5’ end (biotin and digoxigenin, respectively), that enables binding to two specifically modified microscopic beads (covered with streptavidin and antidigoxigenin, respectively). Using a high-resolution dual-trap optical tweezers setup^27^, each of the beads is trapped in a separate optical trap allowing to apply tension on the construct and monitor its extension. Upon introduction of RecBCD in the presence of ATP, the enzyme binds to the blunt end and translocates until it reaches the fork. Then, due to the opposite polarities of RecB and RecD, each of the subunits translocates on an opposite track in an inter-subunit “tug-of-war”, as evidenced by an increase in the tether’s tension. As the force increases beyond a certain level (*F* ≈ 42 − 50 pN), RecBCD dissociates from the construct. Control experiments where the stem’s end was blocked by ligating a short loop or in the absence of ATP showed no translocation activity.

If one track is made significantly shorter than the other, the subunit translocating on the shorter side will reach the bead in a considerable shorter time and, even if futile hydrolysis cycles may take place, translocation on DNA will be impeded. From this point, only the second subunit’s activity will affect the tether’s extension. Hence, by asymmetrically manipulating the tracks’ lengths we can directly measure the activity of individual subunits in the WT RecBCD without the need for mutations. Fig. 2B shows representative traces in three experimental setups to probe the activities of both subunits (left), RecD (middle) and RecB (right), at a wide range of [ATP] and in the force range of 10-15 pN. Velocity curves calculated from all the translocation traces (Supplementary Table 2) indicate that the full enzyme, as well as its subunits, display an apparent hyperbolic dependence on [ATP], with *K*_1/2_ = 200 ± 49 µM for RecD and *K*_1/2_ = 77 ± 24 µM for RecB. The measured *K*_1/2_’s are consistent with previously reported Michaelis-Menten fits for RecBCD ^23, 28^ and RecBC ^10^. Remarkably, both of the *K*_1/2_′*s* values are significantly smaller than the nucleotide concentrations at which the second binding phase was observed (*K*_*w*_ > 280 µμM, Supplementary Table 1). Since we measured similar affinities of RecBCD towards different nucleotides, our results indicate that both subunits’ catalytic sites are bound by ATP at concentrations that are lower than the weak binding phase, and further rule out the possibility of a biphasic curve arising from ATP binding to the two catalytic sites of each of the subunits of RecBCD. Corollary, they further support the identification of the higher affinity sites with RecBCD’s catalytic binding sites, and the lower affinity sites with previously uncharacterized auxiliary binding sites for ATP.

(Of note, the existence of the secondary translocase activity discovered by Lohman and coworkers^10^ implies that the RecB subunit translocates in both the 3’ to 5’ and 5’ to 3’ direction. Hence, while the third construct of Fig. 2B probes the activity of RecB’s *primary* translocase, providing an upper bound for the *K*_D_ of RecB’s catalytic site, in the second construct two types of 5’ to 3’ activities are present: RecD’s translocase and the *secondary* translocase of RecB. Since both activities in RecB are derived from the same ATPase reaction^10^, these experiments set an upper bound on the *K*_D_’s of both RecB and RecD. Together, they lead to the same conclusions as stated above, without considering the secondary translocase activity of RecB).

### ATP binding to the strong and weak binding sites are kinetically separated

Given the distinct location and chemical nature of the additional binding sites, we examined whether they affect also the kinetics of nucleotide binding to RecBCD. Complementary to the equilibrium binding assays, transient kinetics can reveal the reaction pathways of binding sites and the order of binding. Unfortunately, the exceedingly rapid ATP turnover of RecBCD (one catalytic turnover <1 msec) implies that all biochemical transitions along the ATPase cycle occur at very fast rates, making the characterization of the binding kinetics very challenging. However, at 6 °C, when the ATP turnover is significantly reduced (Supplementary Fig. 5) we were able to monitor the transient kinetics of mant-nucleotides binding to RecBCD (Fig. 3A). For simplicity, we show here our analysis for mantADP only, and refer the reader to the Supplementary Information (Supplementary Results, Supplementary Fig. 6 and Supplementary Table 3) for the analysis of additional binding kinetic experiments performed with mant-ATP, and in the presence and absence of DNA.

**Figure 5:**
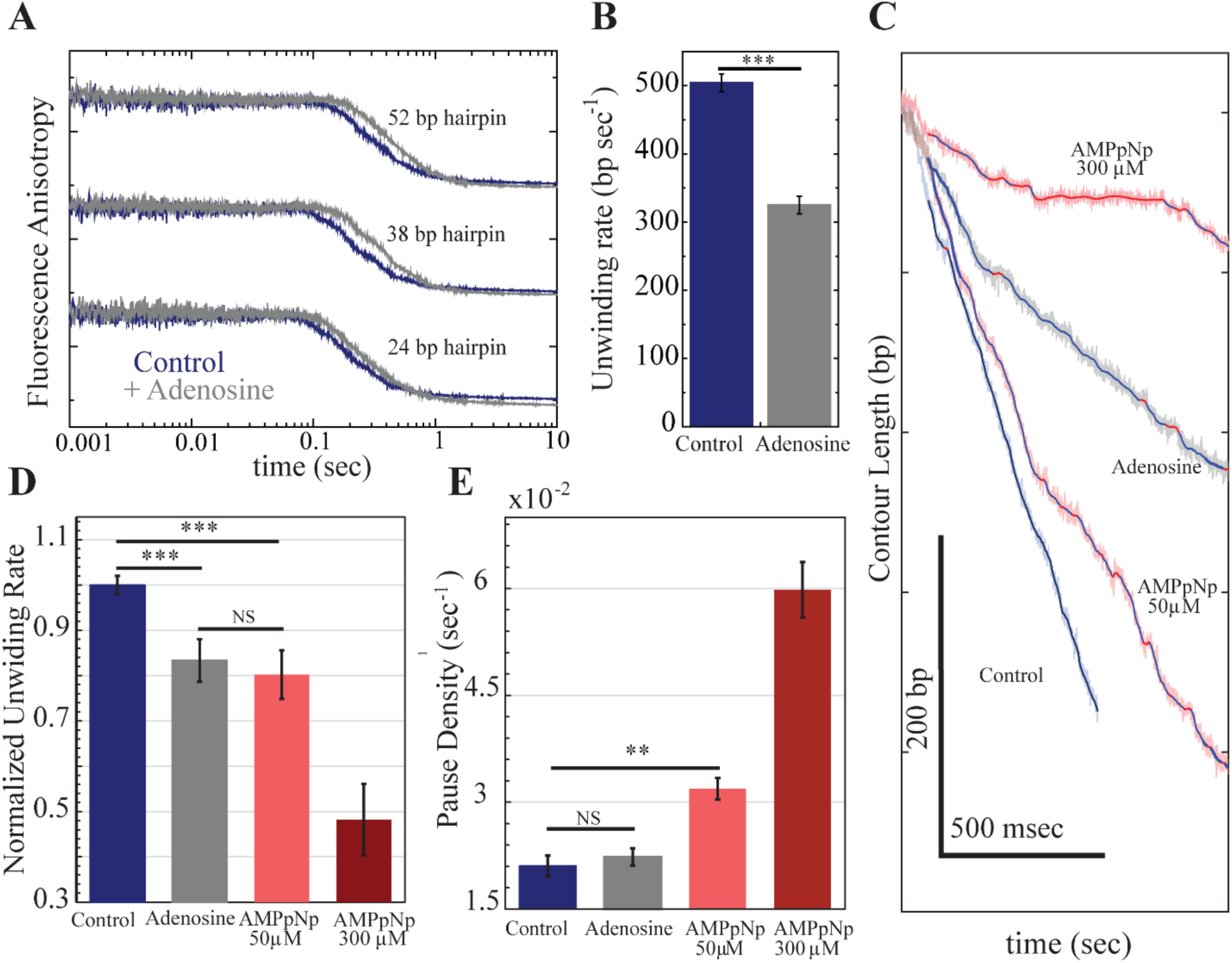
Adenosine slows down RecBCD without halting it. **A.** FA time courses of unwinding reactions for a RecBCD× hpDNA complex (250 nM, post-mixed; top: 52 bp, middle: 38 bp, bottom: 24 bp) at [ATP] = 350 μM and [NaCl]=75mM, in the absence (blue) and the presence (grey) of 2mM adenosine. Time courses were vertically shifted for clarity. **B.** The unwinding rate in the absence and the presence of 2 mM adenosine. Data shown as mean ± s.e.m., n = 15, two sample Student’s t-test, ***P<0.001. **C.** Representative traces of single-molecule unwinding/translocation experiments in the presence of adenosine (gray) and AMPpNp (light red, 50 µM and dark red, 300 µM), at 2 mM ATP. Also shown is a control (blue) in the absence of both adenosine and AMPpNp. Detected pauses are marked in red for all traces. **D.** RecBCD’s normalized translocation rates in the absence and presence of adenosine or AMPpNp. Data shown as mean ± s.e.m., n=34,9,21 and 22 for control, adenosine, 50 µM AMPpNp and 300 µM AMPpNp, respectively. Two sample Student’s t-test, *** P<0.001, NS non-significant. Color coding as in A. **C.** Pause density in the absence and the presence of adenosine and AMPpNp. Data shown as mean ± s.d., n=34, 9, 21 and 22 for normal, adenosine, 50 µM AMPpNp and 300 µM AMPpNp, respectively. Two sample Student’s t-test, *** P<0.001, ** P <0.01, NS non-significant.

**Figure 6:**
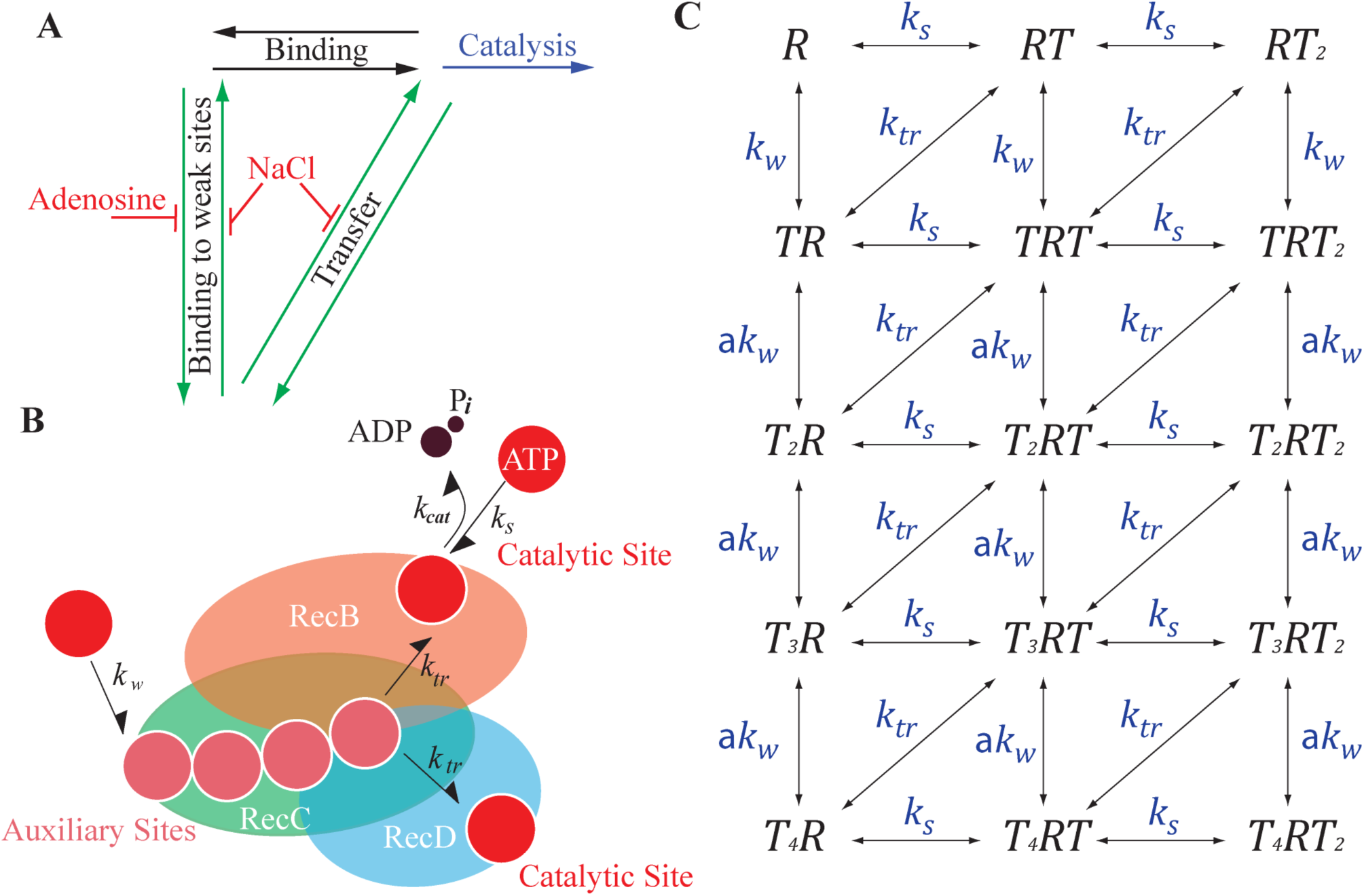
Transfer of ATP from RecBCD’s auxiliary sites to the catalytic ones supports rapid catalysis. **A.** Binding of ATP to RecBCD’s catalytic sites can be achieved through two parallel pathways: directly from the solution (black), or through cooperative binding to the auxiliary sites followed by a transfer step (green). **B.** RecBCD utilizes the auxiliary binding sites as an ATP buffer, facilitating binding to the catalytic sites. The RecBCD enzyme is represented by orange, green and blue ovals for RecB, RecC and RecD, respectively. **C.** Schematic representation of chemical intermediates involved in ATP binding to RecBCD. R represents RecBCD, T represents an ATP molecule. *T*_*n*_*RT*_*m*_, represents *n* ATP molecules bound to the non-catalytic (auxiliary) sites and *m* ATP molecules bound to catalytic sites. Binding to RecBCD catalytic sites is uncooperative with rate *k*_*s*_, binding to the non-catalytic sites, *k*_*w*_, is cooperative (binding to the second, third or fourth non-catalytic sites is faster by the factor *a*). Transfer from the non-catalytic sites to the catalytic sites occurs at a rate *k*_*tr*_. The reverse rates corresponding to each step are not shown, for the sake of clarity.

For all concentrations measured, mantADP binding to RecBCD exhibited two kinetic phases, and can be well described by the sum of two exponential functions (Figs. 3A, B). Interestingly, the observed rates for both the faster and slower phases 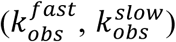 exhibited a hyperbolic concentration dependence (Fig. 3B), indicating a minimal two-step mechanism where the initial binding is followed by an isomerization step that results in a high fluorescence complex. Moreover, the saturation kinetics measured for the fast rate suggests that nucleotides binding to RecBCD takes place through parallel, independent pathways, rather than a single pathway of multiple, sequential binding events ^29^.

To elucidate how the two phases observed in these experiments (‘fast’ and ‘slow’) correlate with the previously characterized binding sites (‘strong’ and ‘weak’), we measured the transient kinetics of binding in the presence of 2 mM adenosine, which was shown above to block the weak sites with no significant effect on the strong ones. Remarkably, a single exponential phase was observed under these conditions (Fig. 3C), with a hyperbolic concentration dependence and similar values to the fast phase observed in the absence of adenosine (Fig. 3D). Hence, we conclude that the fast phase represents binding of ATP to the strong, catalytic sites, while the slow phase represents binding to the weak sites.

### Salt slows RecBCD’s unwinding velocities exclusively at high ATP concentration

To elucidate the functional role of the weak sites on the unwinding activity of RecBCD, we performed a real-time, single turnover, fluorescence anisotropy (FA) unwinding assay, where pre-incubated RecBCD·hpDNA is rapidly mixed with ATP and ssDNA traps for the unwound hpDNA and the dissociated RecBCD (Fig. 4A). The time courses of unwinding reactions exhibited a lag phase followed by a decay in FA (Fig. 4B). We measured the unwinding transients for three substrates of different lengths (24bp, Supplementary Fig. 7A; 38bp, Supplementary Fig. 7B; and 52 bp, Fig. 4B), in two [ATP] regimes (100 and 250 µM), and with varying [NaCl] conditions. The choice of the [ATP] regimes was based on the equilibrium binding curves, and meant to test the effect of the weak binding sites through manipulation of [NaCl]. All unwinding traces were fitted to Eq. 5 (Methods), and the unwinding lag phase duration is shown in Fig. 4C as a function of the substrate length.

The total time unrelated to unwinding, indicated by the intercept of *t*_*lag,uw*_ with the ‘x’-axis, remained constant regardless of [NaCl] and [ATP], indicating that the efficiency of complex formation during the pre-incubation time was not affected in the range of [NaCl] studied. Next, we quantified the slopes of the lag time vs. substrate length curves in Fig. 4C to obtain the unwinding rate (Fig. 4D). Of note, at 75 mM salt, the rates measured for both 100 and 350 µM ATP are consistent to the ones reported in previous works measuring RecBCD’s translocation and unwinding in bulk ^10, 11^ and single molecule assays ^22, 28^. Remarkably, the results show that at 350 µM ATP, when the weak binding sites are ∼ half occupied and hence are sensitive to salt, the unwinding rate decreases as [NaCl] is increased (Fig. 4D and Supplementary Table 4). However, at 100 µM ATP, where the weak binding sites are mostly unoccupied, salt did not have any effect on RecBCD’s unwinding rate. These results strongly support that binding to the weak sites is functionally important in order for RecBCD to achieve its rapid unwinding rate at high [ATP].

### Inhibition of the auxiliary binding sites slows down RecBCD, without inducing pauses

Given the facilitating effect of NaCl on nucleotide binding to the auxiliary sites (Fig. 1B), and the concurrent slowing down of the unwinding velocity (Fig. 4D), one may postulate that ATP binding to the weaker sites may play a negative allosteric function. If that is the case, preventing binding to these sites should increase the unwinding rate. Therefore, we tested their function by the use of adenosine to inhibit ATP binding to these sites during the catalysis of an unwinding reaction. We compared unwinding velocities by time-resolved fluorescence anisotropy at 250 µM ATP in the presence and absence of 2 mM adenosine (Fig. 5A). Surprisingly, adenosine slowed down the unwinding velocities significantly (> 10-fold, Fig. 5B), suggesting a slower mode for unwinding when the weaker binding sites are blocked, and implying a role for these sites, beyond a simple allosteric effect.

To further investigate the underlying mechanism by which adenosine slows down the velocity of RecBCD, we tested whether this is due to a competition with ATP for the catalytic sites. We used our single molecule assay to compare the effect of adenosine with that of AMPpNp. Fig. 5C shows typical translocation traces at 2 mM ATP in the absence and the presence of adenosine (2 mM) or AMPpNp (50 µM and 300 µM). Remarkably, while AMPpNp induces ubiquitous pauses in the traces, an indication of a nonhydrolyzable analog bound at the catalytic site, adenosine slows down RecBCD without introducing measurable pauses. In particular, whereas the average velocity of translocation is comparable for 2 mM adenosine and 50 µM AMPpNp (Fig. 5D), and slower as compared to the ATP-only case, the slowing down is due to different mechanisms: the presence of AMPpNp results in an increase in the pause density, while for adenosine the slowing down occurs without significant pauses as compared to the ATP-only case (Fig. 5E), suggesting that the pause-free, *instantaneous* rate is decreased by preventing binding to the weak sites.

### A kinetic scheme that includes transfer of ATP to the catalytic sites recapitulates the data

Our data indicates that blocking ATP binding to the weaker sites by adenosine lowers the unwinding velocity of RecBCD, therefore suggesting that binding to these sites plays a role in the catalytic cycle. In addition, since salt reduces the unwinding rate at high [ATP], it seems that the ability to dissociate from these sites is important as well, arguing against an allosteric mechanism of catalysis regulation by binding to the weak sites. Hence, we suggest a model in which ATP can bind to the weak sites and transfer, perhaps by diffusion on the surface of the protein, to the catalytic site (Fig. 6A). The premises of the model are: 1) The catalytic sites are non-cooperative (for simplicity, we assume they are identical), 2) There are additional, weak sites that are cooperative (for simplicity, we assume 4 such sites), 3) There is no cooperativity/allostericity *between* the weak and the strong binding sites, 4) Unwinding can occur from any state that includes ATP in at least one of the catalytic sites, and 5) There is a transition of ATP from the non-catalytic sites to the catalytic sites. Our proposed scheme is shown in Figs. 6A, B. In this framework, the fast binding of ATP to the auxiliary sites, followed by its transfer to the catalytic site, can serve as an additional pathway contributing to the productive flux of ATP to the catalytic site.

Based on this scheme, we used thermodynamics to calculate an expression for the total equilibrium ATP occupancy when there is no catalysis, and simulations to find the rates of pre-equilibrium binding and of unwinding activity (Methods). We then globally fitted the model to our measurements of the equilibrium and kinetics of nucleotides binding and the unwinding activity, with a set of kinetic parameters 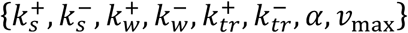, and the assumptions that adenosine blocks binding to the auxiliary sites (i.e. lowers 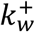) and salt affects the dissociation of ATP (i.e. reduces 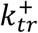 and 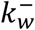). Notably, our global fitting analysis recapitulates all the data (Supplementary Fig. 8 and Supplementary Table 5), and reveals as expected that RecBCD’s additional binding sites are characterized by slow binding and dissociation rates, weak affinity binding to ATP, and sensitivity to adenosine and salt (Supplementary Table 5 and Supplementary Fig. 9). The exchange of ATP across these sites is essential for obtaining the measured velocities for RecBCD. Additionally, analysis of the modeled kinetics confirms that the both the occupancy and the release from those sites are essential for fast binding of ATP to the catalytic sites (Supplementary Fig. 10). In the presence of adenosine, the binding to the catalytic sites is slightly affected by blocking the non-catalytic sites, emphasizing that binding to those sites is critical. Moreover, while blocking the release of ATP from these sites increases their occupancy (due to lower 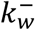 and 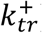), the overall occupancy of the catalytic sites decreases resulting in lower unwinding rates.

It may be considered, that the biphasic nature of the binding isotherms may result in a deviation from Michaelis-Menten kinetics, in contrast with the hyperbolic kinetics measured for unwinding by RecBCD as a function of [ATP], here and by others ^23, 28^. However, it is important to note that although two phases are observed in the equilibrium binding curves, the unwinding curve represents a steady state distribution, not equilibrium. Hence, it is governed by the kinetic rates of ATP flux between the different ATP bound states (Fig. 6C), i.e. on the microscopic kinetic rate constants *k*_*s*_, *k*_*tr*_ and *k*_*w*_, and the inherent competition between the binding pathways. As a result, for certain values of these kinetic parameters (like those found in our global fit), the model results in kinetics that resembles, within experimental uncertainty, a hyperbolic curve. This is shown in Supplementary Fig. 11, were the results from our global model show that the unwinding rate as a function of [ATP] shows a hyperbolic-like dependence, with and without adenosine.

## DISCUSSION

RecBCD initiates the DSB response in *E. coli* by unwinding DNA at a remarkably fast rate, much higher than the rates of other related helicases. In our work, we aimed to elucidate what are the specific characteristics that allow RecBCD to catalyze the unwinding reaction at such high velocities. We reveled that, in addition to the known catalytic sites, RecBCD harbors auxiliary nucleotide-binding sites of functional importance. Equilibrium binding titration curves exhibited a biphasic behavior that cannot be accounted by two catalytic sites only, hence suggesting the existence of additional binding sites. These results were supported by equilibrium dialysis experiments, which showed that RecBCD can bind at least 4 nucleotides, in the presence and absence of DNA. Computationally scanning the surface of the complex revealed potential binding sites for ATP, with important clusters in the RecC subunit. Indeed, binding titration curves with purified RecBC indicated that it binds to mant-ADP also with a biphasic isotherm demonstrating that weak sites are located in the RecC subunit. Binding experiments in the presence of NaCl and adenosine allowed us to show that the chemical nature of the strong and weak binding sites is different. Kinetic experiments showed that ATP binds to the different binding sites independently, and with different kinetic properties: faster for the strong sites and slower for the weak ones. Single molecule experiments probing the translocation activity of the RecB and RecD subunits in the native context of a WT RecBCD complex, showed that both subunits are active at similar [ATP], allowing us to conclude that the strong sites correspond to the known catalytic ones, while the weak ones are previously uncharacterized binding sites for ATP. Finally, we showed that both salt and adenosine slow down the activity, indicating that both association and dissociation of ATP at these sites are required to increase the catalytic rate at high [ATP]. Based on these observations, we proposed a kinetic model where the newly-found auxiliary sites bind ATP and transfer it to the catalytic site. A global fit of the model to our equilibrium binding, pre-steady state binding kinetics and unwinding activity is able to capture all our results.

Remarkably, our biochemical and computational results are in line with previous evidence for the existence of additional ATP binding sites in RecC. In the pioneering work by Julin and Lehman 30 years ago ^30^, using photoaffinity labeling of RecBCD with 8-Azidoadenosine 5’-Triphosphate, the authors reported an apparently non-specific crosslinking of ATP to RecC, indicating the possibility of additional ATP binding site/s. These experiments were limited to 200 μM modified nucleotide concentration and hence could only detect the initial weak binding. However, it is possible that these earlier results provided and indication for the existence of the auxiliary sites characterized in our work.

Our results reveal a new mechanism supporting the rapid turnover of RecBCD, in which the newly-characterized sites serve as a “buffer” of ATP molecules, which can quickly transfer to the catalytic sites upon release of the hydrolysis products resulting from the previous cycle. This mechanism will effectively enhance the flux of ATP to the catalytic site under a specific kinetic premise: The overall flux of ATP molecules from the auxiliary sites to the catalytic ones, which includes ATP binding to the auxiliary sites and its subsequent transfer to the catalytic sites, must be larger than the rate of binding directly from the solution. The global fitting results indicate that the transfer rate is indeed very fast (∼4·10^5^ sec^−1^). However, the rate of binding to the auxiliary sites is lower than binding directly to the catalytic ones, as evidenced also from the transient kinetic experiments. This apparent contradiction is settled by taking into account that the auxiliary sites are highly cooperative. As a result, the rate constant for ‘replenishing’ the buffer is given by *a* · *k*_*w*_ which, since *a*∼18, is larger than *k*_*s*_.

Interestingly, the proposed mechanism, where ATP molecules bind to non-catalytic sites to increase the flux of ATP to the catalytic sites, bears some similarities to the facilitated diffusion mechanism postulated for the diffusional search of a transcription factor for its binding site, where 3D diffusion (i.e. binding from the solution to the specific binding site) is combined with non-specific binding and 1D diffusion on the DNA^31^. Also similar is the enhancement of reaction rates by surface diffusion that follows reversible adsorption of ligands^32, 33^. Common to these models is a reduction in the dimensionality of the diffusional search.

The direct binding pathway will be the predominant one (i.e. there will be no buffering effect) both at high [ATP] (when *k*_*s*_[ATP] ≫ k_tr_), and at very low [ATP] (when [ATP]≪K_w_ and the buffer sites are unoccupied). Hence, the importance of the buffering mechanism described here is in supporting rapid RecBCD activity in the middle range, ∼100-300 μM. This raises the question of its physiological role, as average ATP levels are in the milimolar range. Interestingly, recent works have shown that there is a considerable variability between cells in a bacterial population, with a significant fraction of them exhibiting ATP levels much lower than the average ^34^. The mechanism proposed here can serve to ensure proper function of RecBCD, and hence proper damage repair, across the whole population. Finally, a dramatic decrease in intracellular ATP levels occurs upon exposure of cells to reactive oxygen species, such as those released by the host defense systems when it attempts to eliminate an invading bacterium^35^. Although these species do not directly create DSBs, excessive oxidative stress can lead to DSBs via the conversion of unrepaired single strand breaks. Hence, it is possible that the mechanism proposed here evolved as a defense mechanism, to support the activity of RecBCD at full speed during such damage-rich stress situations.

## Acknowledgments

We thank Dr. Stephen C. Kowalczykowski and Dr. Theetha Pavankumar (University of California, Davis, California) for the RecBCD expression system and their assistance with the purification protocol. We are grateful to Dr. Piero R. Bianco (University at Buffalo, Buffalo, NY) for the RecBC null strain and RecBCD plasmids, and for his excellent advice. This research was supported by The Israel Science Foundation [grants 296/13 to AH, and 1750/12 to AK]; the Marie Curie Career Integration Award [grants 1403705/11 to AH and 293923 to AK], and the Israeli Centers of Research Excellence program (I-CORE, Center no. 1902/12 to A.K).

## Author contributions

RZ and VG performed the experiments. DY and SR prepared and provided experimental materials. OM and AK designed and built the optical tweezers setup. AH designed the ensemble FA unwinding assay for RecBCD. RZ, VG, AK and AH analyzed the data and wrote the paper. AK and AH designed and supervised the research.

## Additional information

Any supplementary information, chemical compound information and source data are available in the online version of the paper. Reprints and permissions information is available online at http://www.nature.com/reprints/index.html. Correspondence and requests for materials should be addressed to AK or AH.

## METHODS

### Reagents and purification of proteins

All chemicals and reagents were the highest purity commercially available. ATP and ADP were purchased from Roche Molecular Biochemicals (Indianapolis, IN, USA). Adenosine 5′-(β,γ-imido)triphosphate (AMPpNp) was purchased from Sigma (St. Louis, MO, USA). A molar equivalent of MgCl_2_ was added to nucleotides immediately before use. Nucleotide concentrations were determined by absorbance using an extinction coefficient ε_259_ of 15,400 M^−1^ cm^−1^. The concentrations of N-methylanthraniloyl (mant) derivatives of ADP, 2’-deoxyADP, ATP, and 2’-deoxyATP (Jena Bioscience, Jena, Germany) were determined using ε_255_ of 23,300 M^−1^ cm^−1^. Unless otherwise specified, all experiments were conducted in RecBCD Buffer (RB: 20 mM MOPS pH 7.4, 2 mM MgCl_2_,1 mM DTT, 0.1 mM EDTA and, unless specified, 75 NaCl). Over-expression and purification of recombinant RecBCD was based on the method described by Roman *et. al.*^1^ All steps of purification were carried out at 4 °C, and contained 20 mM MOPS pH 7.4, 2 mM MgCl_2_,1 mM DTT, 0.1 mM EDTA, 1 mM PMSF, 1 mM Benzamidine and the indicated salt concentration. Four liters of *E.coli* cells expressing RecBCD were lysed using Microfluidizer, followed by centrifugation at 10,000×g. The supernatant was further clarified by centrifugation at 100,000×g and treated with Benzonase for two hrs before initial purification by DEAE chromatography (weak anion exchanger to remove nucleic acids contaminants) using a linear NaCl gradient from 75 mM to 700 mM. RecBCD-containing DEAE fractions were eluted from a Q-sepharose column (strong anion exchanger which highly selects for active RecBCD^2^) using a linear NaCl gradient from 75 mM to 1 M. Fractions containing RecBCD were precipitated using (NH_4_)_2_SO_4_ (45% saturation), and collected by centrifugation at 14,000×g. Precipitated RecBCD was resuspended and loaded onto Superdex 200 equilibrated with RB, as a final step of polishing and elution of RecBCD specifically from the monodisperse peak of the heterotrimer complex of RecBCD (Supplementary Fig. 12A-C). Fractions containing purified RecBCD were concentrated using an Amicon concentrator (50 kDa cutoff), aliquoted and flash frozen in liquid nitrogen before storage at - 80 °C. RecBCD concentration was determined using ε_ex,coeff_. of 4.2×10^5^ M^−1^ cm^−1^ in Guanidine chloride. To ensure RecBCD purity from nucleic acids, only protein fractions with 280/260 nm ratio >1.3 were used. RecBC was expressed from a RecBCD-null strain (V330) transformed with pPB700 (RecB), pPB520 (RecC) and pMS421 (lacI) plasmids and purified similarly to the protocol described by Churchill and coworkers^3^. RecBC concentration was determined using ε_ex,coeff_. of 3.7×10^5^ M^−1^ cm^−1^ (Supplementary Fig. 12C). The RecB^K29Q^ plasmid containing the K29Q mutation was obtained by PCR on the pPB700 plasmid using forward (5’-CTGCCGGCACAGGCCAAACCTTTACGATTG-3’) and reverse (5’-CAATCGTAAAGGTTTGGCCTGTGCCGGCAG-3’) primers, using site directed mutagenesis. The mutated plasmid was confirmed by sequencing, and transformed into V330 strain with pPB520 (RecC) and pMS421 (lacI) plasmids. RecB^K29Q^C purification was performed identically to RecBC (Supplementary Fig. 12C). The purified RecB^K29Q^C exhibited negligible ATPase activity (Supplementary Fig. 3C).

To verify that the biphasic observed behavior of nucleotides binding is intrinsic to RecBCD, and not a result of heterogeneity in the protein preparations, we performed an additional control experiment: Immediately after the Superdex 200 purification step, RecBCD fractions were subjected to MonoQ chromatography and eluted with a NaCl gradient as previously published ^2, 4^. This resulted in one major peak (Supplementary Fig. 13A), that was directly assayed for mantADP equilibrium binding as described below. A biphasic isotherm was observed, essentially identical to the one obtained without the Mono-Q purification step (Supplementary Fig. 13B).

### mant-Nucleotide binding by Förster resonance energy transfer (FRET)

FRET measurements were performed with a PC1 spectrofluorimeter (ISS, Champaign, IL), utilizing excitation and emission monochromators. The observation cell was regulated with a Peltier temperature controller at 25 ± 0.1 °C. All equilibrium binding reactions were performed in a 10 μl Precision cell fluorescence cuvette (Farmingdale, NY, USA), which allows minimal inner filter effects^5^ up to a concentration of ∼ 550 μM mantNucleotides. The experiments were conducted in RB, with varying concentrations of NaCl (75, 150, 200, 300 mM). mant-Nucleotides were titrated with a 1:1 ratio to MgCl_2_. Equilibrium binding reactions of mantNucleotides to RecBCD were measured by FRET between RecBCD intrinsic tryptophan fluorescence (*λ*_ex_ = 280 nm) and bound mant-Nucleotide (fluorescence monitored at 90° through an emission monochromator at *λ*_em_ = 436 nm)^6^. We performed subtractions of background fluorescence of free nucleotides on the observed emission peak.

### Determination of equilibrium binding constants and thermodynamics parameters

Nucleotide binding curves of the fluorescence change as a function of the free ligand concentration were fitted to the sum of two Hill equations (Supplementary Information):

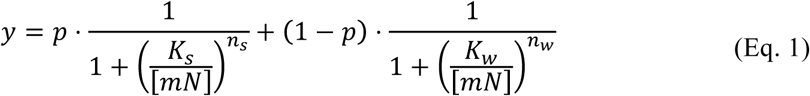

where y is the fraction bound, *mN* is the ligand concentration, *K*_s_ and *K*_w_ are the dissociation constants of the first and second phase, respectively, *n*_*s*_ and *n*_*w*_ are the Hill coefficients of the first and second phase, respectively, and *p* is the partition coefficient (0 ≤ p ≤ 1).

Of note, some uncertainty in the fitting parameters is introduced as a result of a finite overlap between the two phases in the isotherm. Moreover, the relation between the Hill coefficients and the number of sites is only tightly-coupled in the case of infinitely high cooperativity. Hence, the measured coefficients were not used to derive the number of sites.

### Equilibrium dialysis measurements

Equilibrium dialysis measurements were performed with a Fast-Micro-Equilibrium Dialyzer™, with 25 μl chambers, and a Fast Micro-Equilibrium Dialyzer 1 kDa MW cut off membrane™ (Harvard Apparatus, Boston, MA, USA), in RB. Experiments were performed overnight at 4 °C. The concentration of free ligand was determined at the beginning of the experiment by measuring the absorbance at 259 nm (ε = 15.4 M^−1^ cm^−1^, T = 25 °C), and then determined again at the end of incubation from the chamber of the free ligand only. Bound ligand concentration was determined using the mass conservation equation,[N]_i_ = 2[N]_f_ + n[RecBCD], where [N]_i_ is the concentration of the nucleotide at the beginning of the dialysis and [N]_f_ is the concentration at the end of the experiment, from the ligand-only chamber. In the high concentration regime, [RecBCD] was held at 45 μM and [ADP]_i_ or [AMPpNp]_i_ at 1 mM, with a 1:1 ratio to MgCl_2_. For the low concentration regime, we held [RecBCD] at 17 µM and nucleotides at 200 µM. We performed control experiments without RecBCD to estimate the time sufficient to reach equilibrium (overnight incubation) and to validate the level of nucleotides stickiness to the membrane (Supplementary Fig. 14). The total loss of nucleotides across the membrane was less than 0.5%.

### Structural modeling

Binding hot spots on RecBCD (PDB 1W36) were localized using the FTMap computational mapping server (http://ftmap.bu.edu/contact.php), based on 16 small organic molecules as probes (ethanol, isopropanol, isobutanol, acetone, acetaldehyde, dimethyl ether, cyclohexane, ethane, acetonitrile, urea, methylamine, phenol, benzaldehyde, benzene, acetamide, and N,N dimethylformamide)^7^.

### Molecular constructs for single molecule experiments

We generated unwinding/translocation tracks of different lengths similarly to previously described methods^8^. 600 and 4000 bp tracks were obtained using standard PCR reactions (Supplementary Table 6, IDT), nicked using Nt.BbvCI for the Biotin-terminated track and Nb.BbvCI for the Digoxigenin-terminated one (enzymes from New England Biolabs), resulting in complementary 29-nucleotides, flanked with 3 nucleotides (5’-TGC-3’). For the symmetric geometry, the 600 biotin and digoxigenin tracks were mixed at equal molar ratios for DNA annealing, creating a ∼1,200 bp fragment. For the asymmetric geometries, 4000 bp handles were annealed to complementary purchased oligonucleotides with the opposite modification (Supplementary Table 6, HPLC purified, IDT). This resulted in asymmetric handles with 4000 bps and ∼35 nt single stranded DNA on opposite sides. All constructs were ligated to a ∼250 dsDNA stem (‘601’ DNA) generated as previously described ^8^.

### Optical Tweezers

Experiments were performed in a custom-made double-trap optical tweezers apparatus, as previously described^8^. Briefly, the beam from an 852 nm laser (TA PRO, Toptica) was coupled into a polarization-maintaining single-mode optical fiber. The collimated beam out of the fiber was split by a polarizing beam splitter (PBS) into two orthogonal polarizations, each directed into a mirror and combined again with a second BS. One of the mirrors is mounted on a nanometer scale mirror mount (Nano-MTA, Mad City Labs). A X2 telescope expands the beam, and also images the plane of the mirrors into the back focal plane of the focusing microscope objective (Nikon, Plan Apo VC 60X, NA/1.2). Two optical traps are formed at the objective’s focal plane, each by a different polarization, and with a typical stiffness of 0.3-0.5 pN/nm. The light is collected by a second, identical objective, the two polarizations separated by a PBS, and imaged onto two Position Sensitive Detectors (First Sensor). The position of the beads relative to the center of the trap is determined by back focal plane interferometry^9^. Calibration of the setup was done by analysis of the thermal fluctuations of the trapped beads^10^, which were sampled at 100kHz.

### Single-molecule experiments

The complete construct was incubated for 15 min on ice with 0.9 µm polystyrene beads (Spherotech), coated with anti-Digoxigenin (anti-DIG). The reaction was then diluted 1000-fold in RB, with the addition of a 1:1 ratio of Mg·ATP, 0.05 mg/ml BSA, and an ATP regeneration system consisting of 7.5 mM Phosphocreatine and 0.05 mg/ml Creatine phosphokinase. Tether formation was performed *in situ* (inside the experimental chamber) by trapping an anti-DIG bead (bound by DNA) in one trap, trapping a 0.9 µm streptavidin-coated polystyrene beads in the second trap, and bringing the two beads into close proximity to allow binding of the biotin tag in the DNA to the streptavidin in the bead. The laminar flow cell (Lumicks) had 4 channels: streptavidin beads pre-bound to the DNA construct, anti-digoxigenin beads, RB, and RB with the addition of RecBCD. Single DNA tethers were verified in the buffer-only channel and then held at a tension of 5 pN and translocated to the RecBCD channel, until activity was observed as indicated by a rapid decrease in the extension and increase in the force.

### Analysis of single-molecule experiments

Data were digitized at a sampling rate f_s_=2,500 Hz, and saved to a disk. All further processing of the data was done with Matlab (Mathworks). The measured extension was transformed into contour lengths (bps) using the worm-like chain model. Average velocities were calculated using the slope of a linear fit to 100-point smoothed contour lengths in the force ranges of 10-15 pN, where minimal force effect was probed on RecBCD’s rates.

### Steady state ATPase activity

The steady-state ATPase activity of RecBCD (1 nM) was measured by monitoring changes in absorbance at 340 nm using the ATP regenerating NADH coupled assay^11^ at 25 ± 0.1 °C in RB, supplemented with saturating (2 mM) Mg·ATP while varying the [*E.coli*-DNA]. The [*E.coli*-DNA] dependence of the steady state ATPase rate (Supplementary Fig. 5A) was fitted to the quadratic form of the Briggs-Haldane equation:

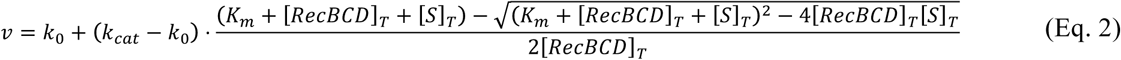

where *k*_0_ is the ATPase rate of RecBCD alone, *k*_*cat*_ is the turnover rate at saturating [S], *K*_m_ is the *apparent* Michaelis constant for substrate activation, [*RecBCD*]_*T*_ is the total RecBCD concentration, and [*S*]_*T*_ is the total concentration of *E.coli*-DNA.

### Pre-steady state kinetic measurements

FRET between RecBCD intrinsic tryptophans (λ_ex_ = 280 nm) and mant-Nucleotides was monitored at 90° through a 400-nm long pass colored-glass filter. For mant-nucleotide binding, photobleaching affected time courses beyond the fitting windows (> 1 sec). Time course traces shown are averages of five to seven shots of 2000-points collected with the instrument in over sampling mode where the intrinsic time constant for data acquisition is ∼ 64 μs. Rapidly mixing the buffer with mant-nucleotide resulted in a linear dependence of the average fluorescence as a function of mant-nucleotides (Supplementary Fig. 15). Thus, data collected was globally fit to the equation 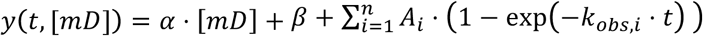, where *α* and *β* describe the linear increase in the initial fluorescence due to increasing mant-ADP ([mD]), and *A*_*i*_ and *k*_*obs,i*_ represent the amplitude and the rate of the measured exponents. Parameter *n* was taken as 2 for normal conditions and as 1 in the presence of adenosine. The dead time of the instrument, determined from the reduction of 2,6-dicholorophenolindophenol with ascorbic acid in absorbance mode, is ∼ 1 msec. Fitting was limited to data beyond the measured dead time. The experiments were conducted in RB, and nucleotides were added in a 1:1 ratio mixture with MgCl_2_.

### DNA substrates for ensemble experiments

DNA oligonucleotides (Supplementary Table 7) were purchased from IDT (Leuven, Belgium) and HPLC purified. The DNA substrates shown in Supplementary Fig. 16 were obtained by folding or hybridization in 20 mM MOPS pH 7.4, 75 mM NaCl, 2 mM MgCl_2_, at 85 °C for 3 minutes followed by slow cooling to room temperature before storage at -20 °C. *E.coli* genomic DNA (Sigma) was digested with EcoRV and SnaBI (NEB) at 37 °C for 4 hrs to create blunt end DNA substrates. DNA concentration was calculated by measuring absorbance at 260 nm and the number of moles of dsDNA was calculated according to *E.coli* genomic restriction map analysis.

### Fluorescence anisotropy monitoring of dsDNA unwinding by RecBCD

Fluorescence anisotropy unwinding time measurements were performed using a T-format excitation and emission module fitted on a SF-61DX2, TGK Scientific (Bradford on von, UK) stopped-flow apparatus thermostatted at 16.0 ± 0.1 °C. The concentrations stated are final after mixing. Samples were excited at λ_ex_ = 492 nm by using vertical polarized light. The emitted vertical and horizontal polarized light was monitored at 90° through a 515 nm long-pass colored glass filter. G-factor for correction of the differences in gain between the dual photomultiplier tube detectors was calculated as described by the instrument manufacturer. Data analysis of the time resolved change in FA was performed according to Henn *et al* ^12^. Equilibrated mixtures of 1.1:1 complex RecBCD·hpDNA (Supplementary Fig. 17) at 250 nM were rapidly mixed with equal volumes of 200 μM or 700 μM Mg·ATP (pre-mixing) and varying [NaCl]. In addition, the ATP solution contained 20 μM of nonspecific ssDNA (25 mer) as a trap for RecBCD, and 5 μM of a non-fluorescent oligo (similar to #2,4,6 in Supplementary Table 7) as a trap for the released hairpin substrate after unwinding. All unwinding reactions were performed at 16 °C to allow sufficient time to monitor the unwinding lag phase. Shown time courses are averages of at least 13-15 transients. The time courses were fitted according to the following function:

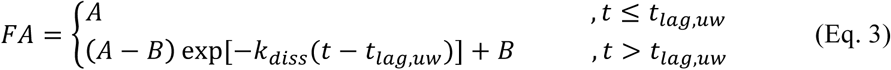

where *A, B* are the initial and final anisotropy values, *t*_*lag,uw*_ is the lag time corresponding to the unwinding duration, and *k*_*diss*_ is the dissociation constant of the oligo from the complex upon release. The fitting was done by minimizing the sum of the squared errors over the parameters {*A, B, k*_*diss*_, *t*_*lag*_} using the Nelder Mead Simplex method in MATLAB. To verify our assay, we show that the lag phase anisotropy corresponds to that of RecBCD bound to the hairpin and the final value of anisotropy is that of ssDNA (Supplementary Fig. 17).

### Detection of pauses in single molecule experiments

Pausing analysis was done by applying a Chung-Kennedy nonlinear adaptive filter^13^, on the contour length *vs*. time data, with windows of size 25, 50 and 100 points and equal weights (Supplementary Fig. 18). The filtered data was processed with a pause detection threshold-based algorithm: peaks of the histogram of the smoothed contour dwell points including more than 50 points were suspected as pauses. To rule out false positives, we applied a series of thresholds on the pause durations (minimal pause 0.002 sec), translocation times between pauses (minimal translocation time 0.001 sec), and contour change between pauses (minimal contour difference 5 bp). Pause density was calculated as the ratio between the total number of pauses and the total contour length of each trace.

### Global fitting to the model

To perform a global fitting to the scheme in Fig. 6C, we calculated its binding partition function or binding polynomial^14^, as:

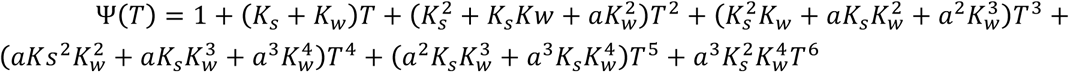

where *K*_*s*_, *K*_*w*_ denote the binding association constants 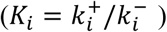 for the strong and weak sites, respectively, *a* the cooperativity constant, and T the ATP concentration. The association constants were calculated as 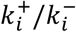 for the strong and weak sites. The fraction of ATP bound is then given by 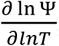, to which our data was fitted. Simulations of mant-nucleotide binding kinetics were generated by numerically solving the system of differential equations describing the the model for up to 30 msec. The time-course of each of the bound states was weighted by the number of bound molecules and the overall weighted sum was fitted to double exponentials. Simulations of DNA unwinding by RecBCD were generated by assuming a direct proportionality with hydrolysis, and numerically solving the resulting the system of differential equations resulting from the model, including the catalysis steps, up to 30 sec. A linear fit to these results was used to calculate the unwinding rates. The sum of the normalized squared error for all experiments (in Supplementary Fig. 7), adjusted for their temperature using the temperature dependent ATPase rates in Supplementary Fig. 4B, was minimized using the global search tool (scatter search algorithm with constrained optimization, interior point) in MATLAB for the parameter set describing: 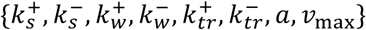, where 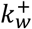 was taken as a non-negative linear decreasing function of adenosine 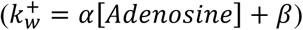, and 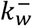 and 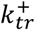 were taken as non-negative-linear decreasing functions of [NaCl] (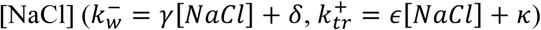 resulting in a global fitting of 11 parameters.

The number of weak binding sites included in the model was based on the equilibrium binding curves: We developed the equations of kinetic schemes similar to the ones in Fig. 6C, where the number of catalytic and non-catalytic sites were varied. For all these cases, the model was globally fitted to the data, and the resulting (calculated) binding curves were compared to the experimental data. We found that four auxiliary sites, in addition to the two catalytic sites, produced the best fitting to our measured data. Increasing the number of weak sites reduces the amplitudes of the catalytic sites, whilst schemes with a smaller number of sites fail to capture the cooperatively measured.

To test the robustness of the model, we applied a bootstrap Monte Carlo method ^15^. We generated 100 random data sets, drawing each parameter from a distribution with a mean equal to the experimentally measured value, and a width equal to the experimental uncertainty. We then fitted each data set to the model using a global search algorithm, resulting in 100 sets of fitting parameters. Histograms of each of the fitted parameters were then fit to Gaussian functions, as shown in Supplementary Fig. 20. This analysis shows that the model robustly converges to a global minimum.

### Data availability

The datasets generated and analyzed during the current study are available from the corresponding authors on reasonable request.

### Code availability

The custom computer code used for the analysis of the results reported here is available from the corresponding authors on reasonable request.

