## Supplementary Materials for "Auxiliary ATP binding sites power rapid unwinding by RecBCD"

### SUPPLEMENTARY RESULTS

#### Nucleotide binding to RecBCD cannot be accounted by the two catalytic sites

Binding of mant-nucleotides to RecBCD and RecBCD·DNA measured by FRET exhibited a biphasic binding pattern. Since the RecBCD complex possesses two well defined catalytic nucleotide-binding sites, in RecB and RecD, one might postulate that these two binding phases reflect binding of nucleotides to each one of these catalytic sites, or that a cooperative model of two binding sites can give rise to such pattern. However, here we show that the biphasic nature of the binding isotherm cannot be accounted for by two binding sites. In what follows we analyze the possible cases involving two binding sites, and show that none of them can give rise to a biphasic curve.

*Model 1: Two independent binding sites.* In this case, each site represents binding of a nucleotide to a catalytic site and the sites are non-cooperative in their binding. It is well known that binding of a single ligand to a single site gives rise to a hyperbolic pattern. Hence, the overall occupancy, $y$, can be expressed as a weighted sum of two hyperbolas, i.e.

$$y = p\frac{x}{x + K_1} + (1 - p)\frac{x}{x + K_2} \qquad \text{(Supplementary Eq. 1)}$$

where $x$ is the ligand concentration and $K_1$ and $K_2$ the dissociation constants of the binding sites. The derivative of this curve,

$$\frac{\partial}{\partial x}y = \frac{pK_1}{(x + K_1)^2} + \frac{(1 - p)K_2}{(x + K_2)^2} \qquad \text{(Supplementary Eq. 2)}$$

is always positive, and decreases with increasing concentrations. Therefore, $y$ will always have a hyperbolic pattern regardless of the values of $K_1$ and $K_2$. Simply put, the sum of two concave hyperbolas is also a concave function (Supplementary Fig. 2A). Since for a biphasic pattern, the derivative of the curve should decrease in the first phase and then increase in the second, this model cannot give rise to a biphasic pattern.

*Model 2: A Hill model for two cooperative binding sites.* The Hill equation,

$$y = \frac{x^n}{x^n + K^n} \qquad \text{(Supplementary Eq. 3)}$$

is the most common way to describe cooperativity. Two cooperative sites on a macromolecule will result in a Hill coefficient that is non-unity. For positive cooperativity, the Hill coefficient is greater than unity resulting in a sigmoidal pattern which cannot explain the data (Supplementary Fig. 2B). For negative cooperativity, the Hill coefficient is lower than unity resulting in a derivative,

$$\frac{\partial}{\partial x}y = \frac{nK^n x^{n-1}}{(x^n + K^n)^2} \qquad \text{(Supplementary Eq. 4)}$$

that is always positive and decreasing with increasing ligand concentration, resulting in a hyperbolic-like curve (Supplementary Fig. 2C). Hence, this model cannot explain our data.

Thus, two sites are insufficient to give rise to a biphasic binding pattern. Following this argument, one can decompose the binding curve into two phases, one hyperbolic with high affinity and one sigmoidal with weak binding. The decomposition is shown in Supplementary Fig. 2D. The first hyperbolic phase could be due to a single site, or multiple loosely coupled binding sites. The second binding phase represents a sigmoidal pattern, necessitating additional sites (at least two, and cooperative) that are distinct from the site/s giving rise to the first phase. Assuming that the affinities of the weak and strong sites are well separated, we phenomenologically describe the total occupancy by the weighted sum of two Hill equations:

$$y = p\frac{x^{n_1}}{x^{n_1} + K_s^{n_1}} + (1-p)\frac{x^{n_2}}{x^{n_2} + K_w^{n_2}} \qquad \text{(Supplementary Eq. 5)}$$

where $(K_s, K_w)$ are the microscopic dissociation constants, and $(n_\mathrm{s}, n_\mathrm{w})$ the Hill coefficients of the strong and weak phases, respectively. As shown in Supplementary Fig. 2D, this model can result in a biphasic pattern. Figs. 1A, B and D show that it fits our data well.

**mant-Nucleotides binding kinetics detailed analysis**

Time courses of mantADP (Fig. 3A) and mantATP (Supplementary Fig. 6) binding to RecBCD displayed a double-exponential pattern. The fast phase observed rate $\left(k_{obs}^{fast}\right)$ of mantATP and mantADP binding to RecBCD and RecBCD·DNA complexes exhibited a hyperbolic concentration dependence on mant-nucleotide ($mN$) concentration (Fig. 3B and Supplementary Fig. 6). Hence, it is modeled by a reaction mechanism where a rapid equilibrium step is followed by an isomerization step to a high fluorescence complex $RecBCD{\cdot}mN^*$, as follows:

$$RecBCD + mN \xleftrightarrow{K_1} RecBCD \cdot mN \xleftrightarrow{k_{\pm 2}} RecBCD \cdot mN^*$$

This two-step binding reaction predicts that the observed rate constants for $mN$ binding should obey a rectangular hyperbola according to:

$$k_{obs} = \frac{k_{+2}[mN]}{K_1 + [mN]} + k_{-2} \qquad \text{(Supplementary Eq. 6)}$$

where $K_1$ is a measure of the strength of the collision complex between RecBCD and $mN$, and $k_{\pm 2}$ are the on (+) and off (-) rates of isomerization. Supplementary Table 2 summarizes the rate constants of mant-nucleotide binding to RecBCD and RecBCD·DNA complexes. The kinetics of both mantATP and mantADP binding to RecBCD are extremely fast with a relatively tight collision complex: $K_1$ is in the ranges of tenths of μM in all cases. Additionally, the overall apparent second order binding constants, $k_{+2}/K_1$, are an order of magnitude faster than many other molecular motors studied [1,2]. For comparison, it is ~30 fold faster than Rep; a SF1 DNA helicase [3]. Remarkably, the maximum rate of the observed isomerization is faster than 1000 $s^{-1}$ at 6 °C supporting rapid kinetics of nucleotide binding to RecBCD and RecBCD·DNA complexes. Binding of mantATP to RecBCD or RecBCD·DNA is irreversible, with an intercept being indistinguishable from the origin ($k_{-2} \approx 0$).

The binding kinetics for mantATP and mantADP, in the absence or presence of a DNA substrate, is overall very similar in its mechanism and rates (Fig. 3, Supplementary Fig. 6 and Supplementary Table 2). This may suggest that RecBCD has an overall weak selectivity for the type of nucleotide. The structural implication may be that RecBCD nucleotide binding sites are highly accommodating and translating almost every collision complex to a productive, high affinity state which may promote rapid catalysis and high turnover.

### SUPPLEMENTARY TABLES

**Supplementary Table 1**: Results of fitting the equilibrium binding data to the sum of two Hill equations

| Complex | [NaCl] (mM) | $K_s$(µM) | $n_s$ | $K_w$ (µM) | $n_w$ | $p$ |
|---|---|---|---|---|---|---|
| RecBCD·mMp | 75 | 52 ± 8 | 0.99 ± 0.07 | 287 ± 12 | 12 ± 5 | 0.60 ± 0.04 |
| RecBCD·mD | 75 | 13 ± 3 | 0.95 ± 0.07 | 322 ± 7 | 6.2 ± 0.6 | 0.45 ± 0.03 |
| RecBCD·mD | 200 | 34 ± 5 | 0.8 ± 0.2 | 336 ± 11 | 8 ± 3 | 0.5 ± 0.10 |
| RecBCD·mD | 300 | 29 ± 7 | 1.1 ± 0.3 | 279 ± 11 | 18 ± 6 | 0.5 ± 0.10 |
| RecBC | 75 | 33 ± 10 | 1.4 ± 0.7 | 300 ± 42 | 5 ± 2 | 0.4 ± 0.10 |
| $RecB^{K29Q}C$ | 75 | 23 ± 15 | 1.6 ± 1.2 | 318 ± 12 | 5 ± 1 | 0.1 ± 0.05 |

mD: mantADP (mD); mMp: mantAMP-pNp; $K_s$, $K_w$: Dissociation constants for the strong and weak phases, respectively.

**Supplementary Table 2:** Number of traces in single-molecule unwinding experiments

| **ATP (µM)** | **20** | **100** | **200** | **350** | **500** | **1000** | **2000** |
|---|---|---|---|---|---|---|---|
| **RecBCD** | 10 | 30 | 8 | 25 | 10 | 29 | 34 |
| **RecD** | 5 | 16 | 18 | 28 | 48 | 24 | 52 |
| **RecB** | 18 | 13 | 30 | 26 | 30 | 29 | 52 |

**Supplementary Table 3**: Summary of results from the transient kinetics experiments

| Complex | $k_{obs}^{fast}$ | | $k_{obs}^{slow}$ | |
|---|---|---|---|---|
| | $K_1$(µM) | $A(\text{sec}^{-1})$ | $K_1(\mu\text{M})$ | $A(\text{sec}^{-1})$ |
| ADP | 12 ± 3 | 1160 ± 120 | 10 ± 14 | 134 ± 24 |
| ADP + adenosine | 26 ± 12 | 1260 ± 130 | - | - |
| ATP | 13 ± 3 | 870 ± 30 | 235 ±16 | 390 ± 24 |
| ADP + DNA | 11 ± 6 | 1080 ± 60 | 11 ± 4 | 42 ± 5 |
| ATP + DNA | 38 ± 4 | 1160 ± 70 | 10 ± 6 | 125 ± 70 |

**Supplementary Table 4: Salt dependent unwinding rates.**

| | High ATP | | Low ATP | |
|---|---|---|---|---|
| **[NaCl] (mM)** | **Unwinding rate (bp sec$^{-1}$)** | **p-value[1]** | **Unwinding rate (bp sec$^{-1}$)** | **p-value[1]** |
| **75** | 504 ± 13 | } <0.001 | 130 ± 5 | } NS |
| **150** | 475 ± 21 | } <0.05 | 128 ± 7 | } NS |
| **200** | 430± 12 | } <0.001 | 123± 8 | } NS |
| **300** | 378 ± 10 | | 118 ± 8 | |

[1]Two Tailed, Two sample student t-test

**Supplementary Table 5**: Summary of results from the global fit to the model

| **Kinetic Parameter** | **Value** | |
|---|---|---|
| $k_s^+ (\mu M^{-1}\ \mathrm{sec}^{-1})$ | 56 ± 3 | |
| $k_s^- (\mathrm{sec}^{-1})$ | 678 ± 1 | |
| $k_w^+ (\mu M^{-1}\ \mathrm{sec}^{-1})$ | $\alpha\ (\mu M^{-1}\ \mathrm{sec}^{-1}\ mM^{-1})$ | -4 ± 1 |
| | $\beta\ (\mu M^{-1}\ \mathrm{sec}^{-1})$ | 10 ± 2 |
| $k_w^- (\mathrm{sec}^{-1})$ | $\gamma\ (\mathrm{sec}^{-1}\ mM^{-1})$ | -53 ± 3 |
| | $\delta\ (\mathrm{sec}^{-1})$ | 36,500 ± 29 |
| $k_{tr}^+ (\mathrm{sec}^{-1})$ | $\epsilon\ (\mathrm{sec}^{-1}\ mM^{-1})$ | -487 ± 2 |
| | $\kappa\ (\mathrm{sec}^{-1})$ | 178,000 ± 1,600 |
| $k_{tr}^- (\mathrm{sec}^{-1})$ | 35,400 ± 1,100 | |
| $a$ | 19 ± 1 | |
| $v_{max} (\mathrm{bp\ sec}^{-1})$ | 1,250 ± 4 | |

**Supplementary Table 6:** Oligonucleotides used for building DNA substrates for Optical Tweezers experiments

| **Construct Name** | **Oligonucleotide sequence (5'→3')** |
|---|---|
| Biotin 600 bp track | /5'Biotin/-GCTTTAATGCGGTAGTTTATCA |
| | GCAGCATTAGGAAGCAGCCCAGGCATTAGGAAG CAGCCCAG |
| Dig 600 bp track | /5' Phosphate/- GCATTAGGAAGCAGCCCAGGCTTTATTGCGGTAG TTTATCA |
| | /5'digoxigenin/-GCATTAGGAAGCAGCCCAG |
| Biotin 4000 bp track | /5'Biotin/-GCTTTAATGCGGTAGTTTATCA |
| | GCACTACGCCTCAGCTTGCCCCTCAGCGATGACCTCAG CATTCCCTTTTTTGCGGCATT |
| Dig 4000 bp track | /5'Phosphate/- CTACGCCTCAGCTTGCCCCTCAGCGATGACCTCAGCGT CACTGGTCCCG |
| | /5'digoxigenin/-GCATTAGGAAGCAGCCCAG |
| Biotin short track | /5'Biotin/AACCACCAACCAACAACCACCCAAACCCAAA CCCAAGGTCATCGCTGAGGGGC AAGCTGAGGCGTAGTGC |
| Dig short track | /5'Phasphate/CTACGCCTCAGCTTGCCCCTCAGCGATGAC CAAACCACCAACCAACAACCACCCAAACCCAAACCCA CAC/3'digoxigenin / |

**Supplementary Table 7**: DNA substrates used for the ensemble experiments

| # | Sequence Description | Sequence 5' – 3' | nts |
|---|---|---|---|
| 1 | Stopped flow 24 bp | GGAAGAGAGGAAGAGAGGAAGAGAGGAAGA GGGAGGGAGGGAGGGTTTTCCCTCCCTCCCTC CCTCTTCC | 70 |
| 2 | Stopped flow 24 bp FAM | FAM-TCTCTTCCTCTCTTCCTCTCTTCC | 24 |
| 3 | Stopped flow 38 bp | AAGAGAGGAAGAGGAAGAGAGGAAGAGAGGAAG AGAGGAAGAGGGAGGGATTTTCCCTCCCTCTT | 66 |
| 4 | Stopped flow 38 bp FAM | FAM- CCTCTCTTCCTCTCTTCCTCTCTTCCTCTTCCTCTCTT | 38 |
| 5 | Stopped flow 52 bp | GAGAGGAAGAGAGGAAGAGAGGAAGAGGAAGAG AGGAAGAGAGGAAGAGAGGAAGAGGGAGGGATTT TTCCCTCCCTCTT | 80 |
| 6 | Stopped flow 52 bp FAM | FAM - CCTCTCTTCCTCTCTTCCTCTCTTCCTCTTCCTCTCTT CCTCTCTTCCTCTC | 52 |
| 7 | hpDNA | CATGTGACTCGTTACCTGAGTTTTTACTCAGGTAAC GAGTCACATG | 46 |
| 8 | 5' ohDNA | TTTTTTTTTTCCATGGCTCCTGAGCTAGCTGCAGCTTTT GCTGCAGCTAGCTCAGGAGCCATGG | 64 |

FAM: 6-fluorescein amidi

**SUPPLEMENTARY FIGURES**

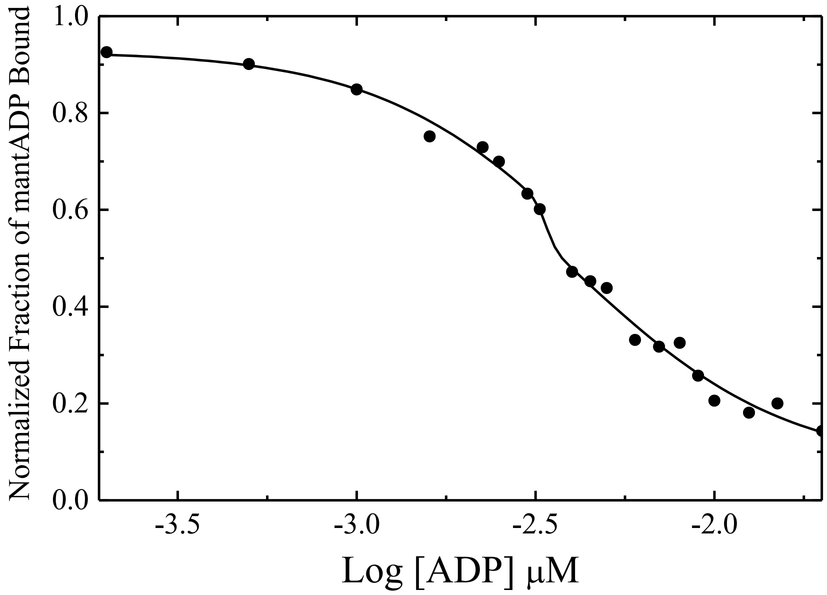


**Supplementary Figure 1: Equilibrium binding competition of unmodified nucleotides to RecBCD·mantADP complexes.** Pre-bound mantADP (600 μM) was competed with titrated unlabeled ADP. MantADP dissociation curve display a biphasic pattern: both nucleotide-binding components, the lower and the higher affinity regimes, are well competed off with the unlabeled ADP.

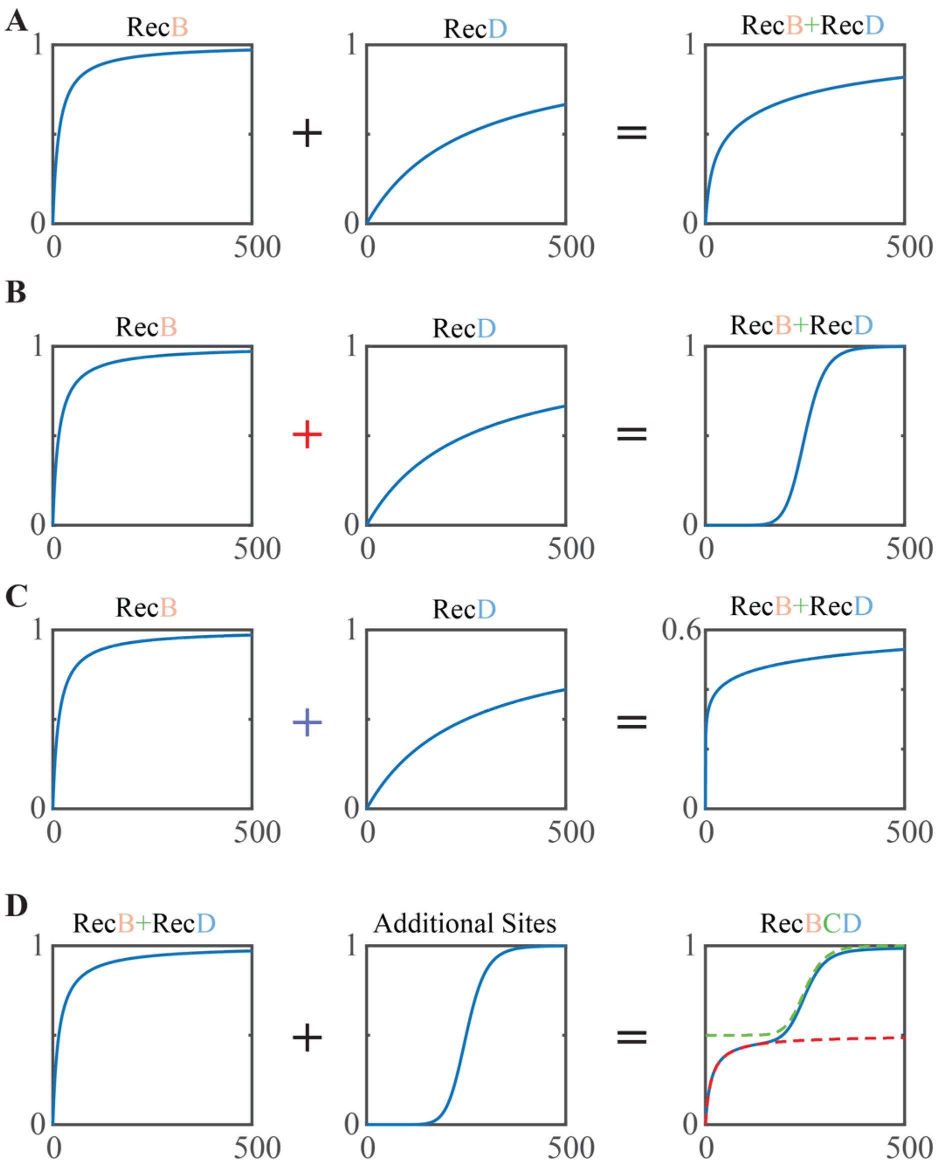


**Supplementary Figure 2: Models for equilibrium nucleotide binding to RecBCD. A.** Nucleotide binding to RecBCD as a result of two binding sites with different affinities. **B.** Nucleotide binding to RecBCD as a result of binding among two sites (catalytic) with positive cooperativity (red plus sign). **C.** Nucleotide binding to RecBCD as a result of binding among two sites (catalytic) with negative cooperativity (blue plus sign). **D.** Nucleotide binding to RecBCD in the presence of non-cooperative binding to the catalytic sites and cooperative non-catalytic sites with lower affinity. The decomposition for the catalytic (red) and additional (green) sites is shown in dashed lines.

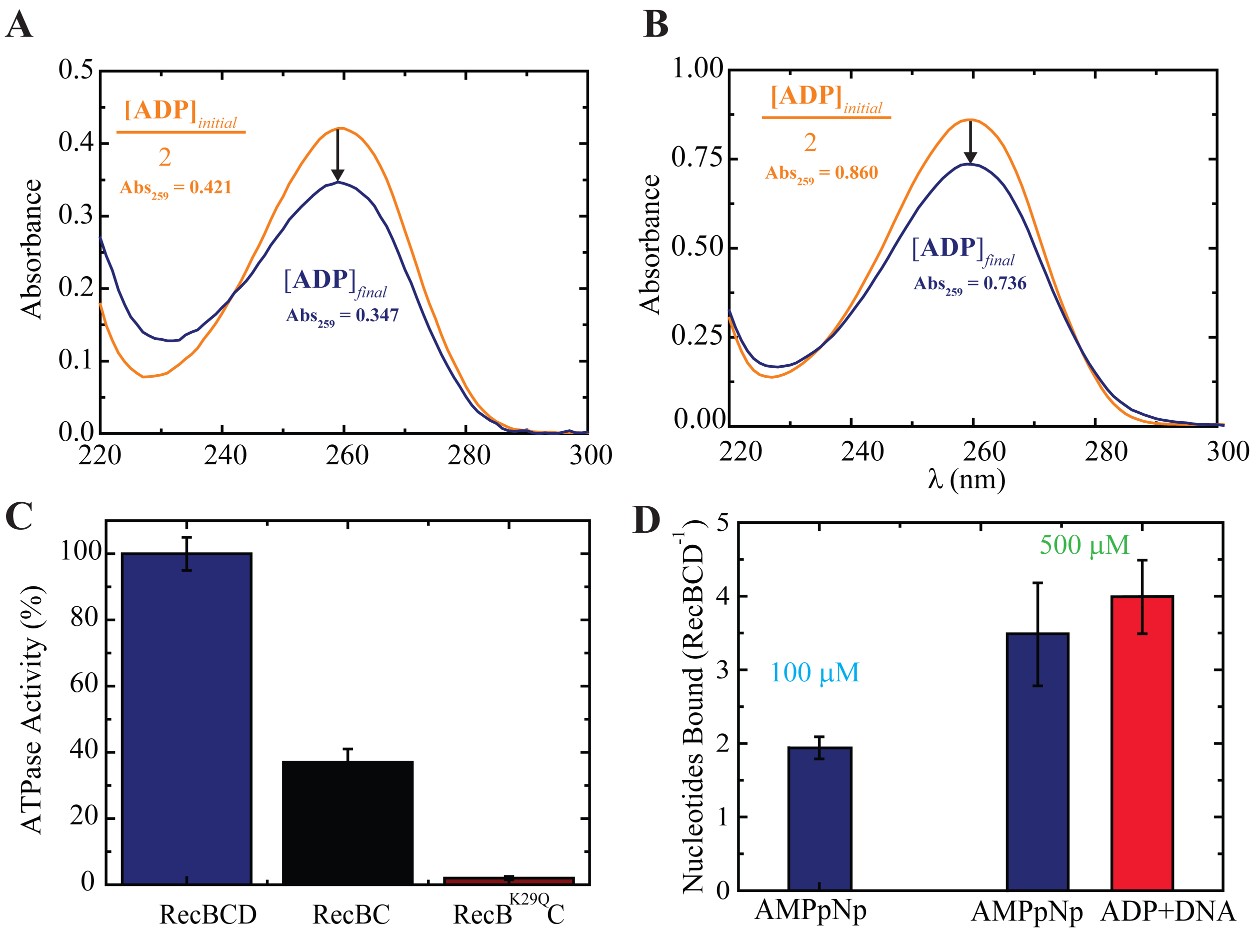


**Supplementary Figure 3: Equilibrium dialysis experiments. A.** Absorbance spectra of 4x diluted ADP pre- (orange, divided by a factor of two) and post- (blue) equilibrium dialysis with 17 μM RecBCD. The net change in the absorption indicates ~2 nucleotide binding sites per RecBCD. Data shown as mean, n = 2. **B.** Absorbance spectra of 10x diluted ADP pre- (orange, divided by factor of two) and post- (blue) equilibrium dialysis with 45 μM RecBCD. The net change in the absorption indicates ~4 nucleotide binding sites per RecBCD. Data shown as mean, n = 4. **C.** Relative ATPase activity of RecBCD, RecBC and the catalytically deficient mutant $RecB^{K29Q}CD$. **D.** Number of nucleotides (blue: AMPpNp, green: ADP) bound to RecBCD (blue) and pre-incubated RecBCD·DNA (green), measured and analyzed as in Fig. 1.

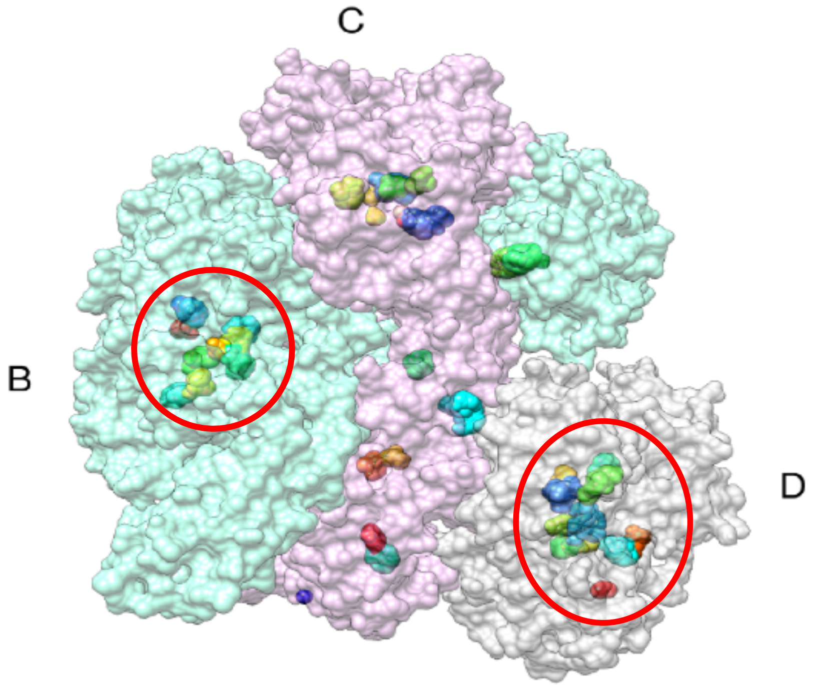


**Supplementary Figure 4: FTMap computation of RecBCD (PDB 1w36).** RecBCD is presented in a space filling model, with chains colored turquoise (RecB), pink (RecC) and gray (RecD). Bound FTmap probes are shown in different colors. The canonical ATP binding sites (Walker A and B) are circled in red.

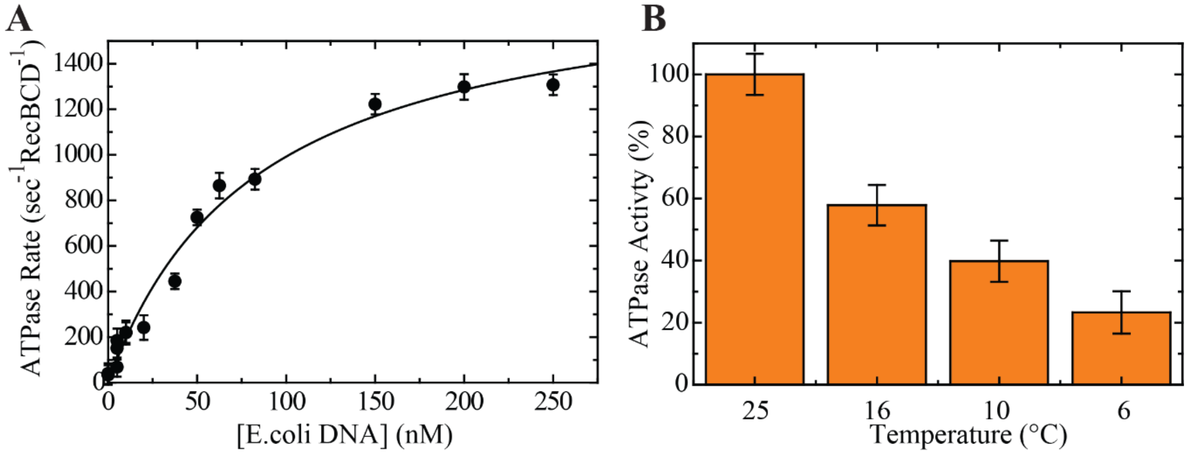


**Supplementary Figure 5: Steady-state ATPase activity of RecBCD with *E.coli* DNA. A.** *E.coli* DNA concentration-dependence of RecBCD steady state ATP turnover velocity (ATPase activity). The solid lines through the data points are the best fits to a quadratic form of the Briggs-Haldane equation (Eq. 2, Methods). The steady state parameters are $K_M$ = 90.9 ± 18.3 nM and $k_{cat}$ = 1,862 ± 143.1 $s^{-1}$ $RecBCD^{-1}$. [*E.coli*-DNA] refers to the concentration of blunt ends in digested *E.coli* chromosomal DNA. Data shown as mean ± s.e.m., n=3. **B.** Temperature dependence of RecBCD ATPase activity, showed as a percent of the mean activity at 25 ºC. Data shown as mean ± s.e.m., n = 3.

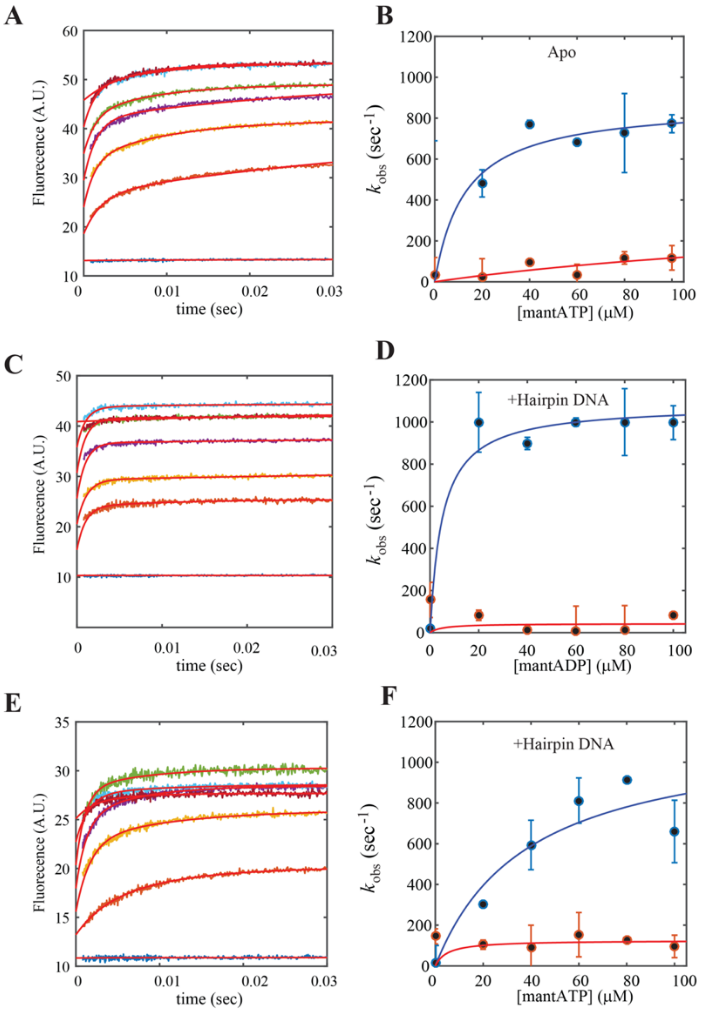


**Supplementary Figure 6: Transient kinetics of mant-nucleotides binding to RecBCD in the absence and presence of DNA. A.** Time courses of mant-ATP binding upon rapid mixing of RecBCD (2 μM, post-mixing) with mantATP (0-100 μM, lower to upper, respectively). The red lines through the data are the best global fit to a double exponential function (Methods). **B.** Dependence of $k_{obs}^{fast}$ (Blue) and $k_{obs}^{slow}$ (Orange) on [mantATP], both displaying hyperbolic dependencies. Data shown as mean ± s.e.m., n = 6-7. The solid line through the data points are best hyperbolic fits. **C.** Time courses of mantADP binding upon rapid mixing of RecBCD·HairpinDNA (2 μM, 1:1.1 ratio, post-mixing) with mantADP (0-100 μM, lower to upper, respectively). The red lines through the data are the best global fit to a double exponential function (Methods). **D.** Dependence of $k_{obs}^{fast}$ (Blue) and $k_{obs}^{slow}$ (Orange) on [mantADP], both displaying hyperbolic dependencies. Data shown as mean ± s.e.m., n = 6-7. The solid line through the data points are best hyperbolic fits. **E.** Time courses of mantATP binding upon rapid mixing of RecBCD·HairpinDNA (2 μM,1:1.1 ratio, post-mixing) with mantATP (0-100 μM, lower to upper, respectively). The red lines through the data are the best global fit to a double exponential function (Methods). **F.** Dependence of $k_{obs}^{fast}$ (Blue) and $k_{obs}^{slow}$ (Orange) on [mantATP], both displaying hyperbolic dependencies. Data shown as mean ± s.e.m., n = 6-7. The solid line through the data points are best hyperbolic fits.

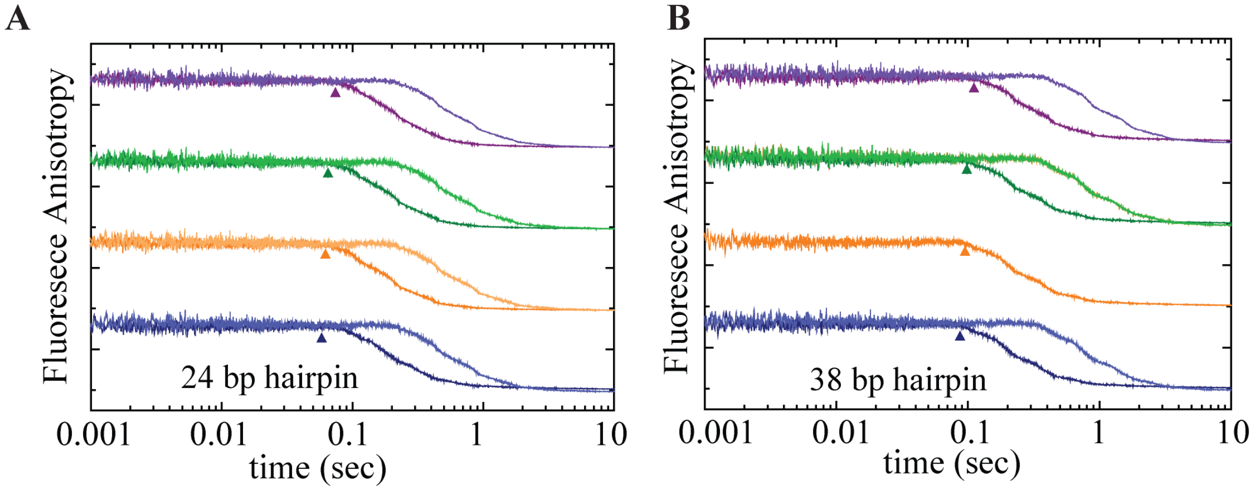


**Supplementary Figure 7: Unwinding time courses of 24 bp DNA and 38 bp DNA. A. and B.** FA time courses of unwinding reactions of a post-mixed 250 nM RecBCD·hpDNA (24bp **A,** and 38bp **B**) with [ATP] = 350 µM (strong colors) and [ATP] = 100 µM (light colors). Color coding as in Fig. 3A.

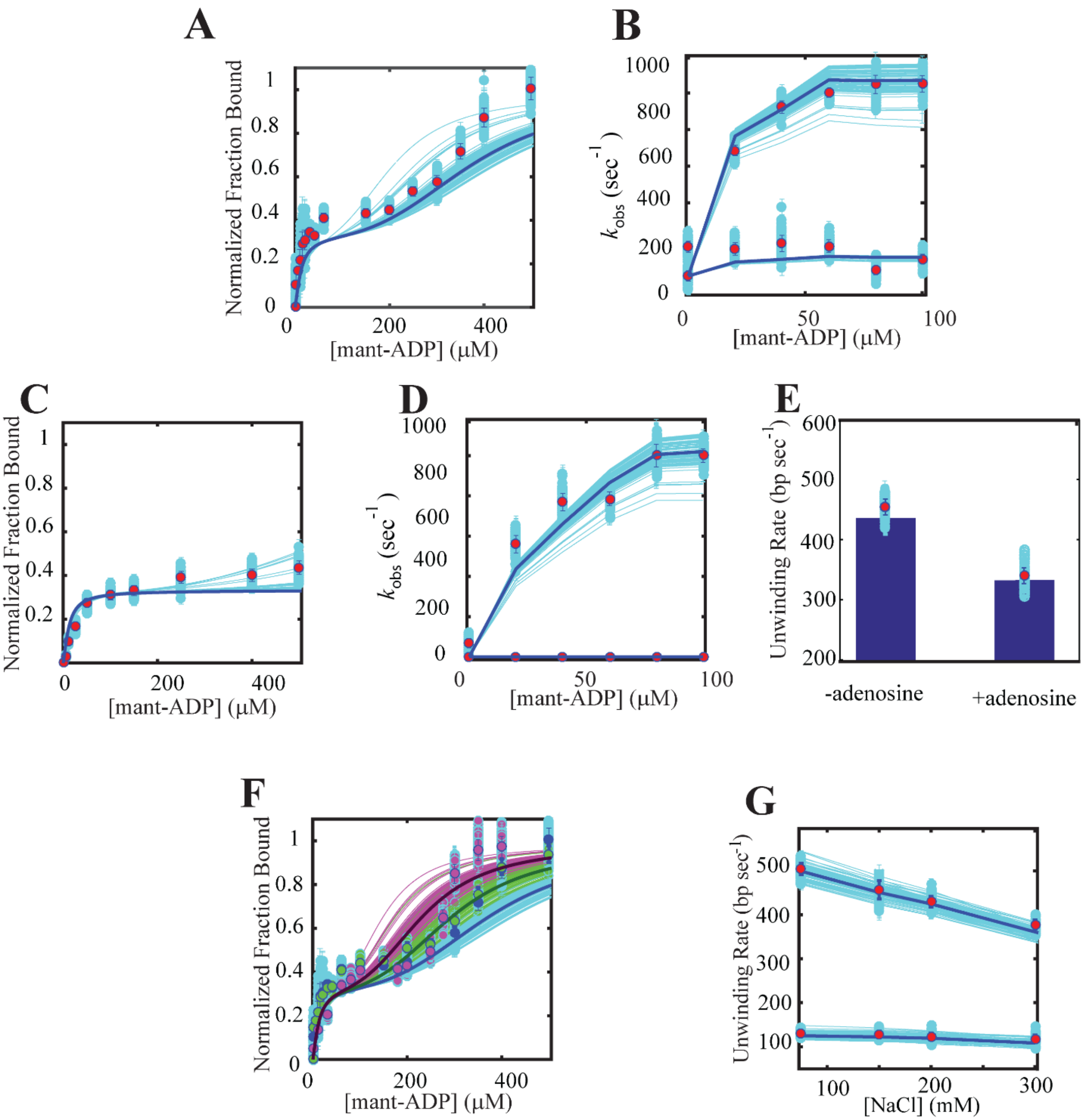


**Supplementary Figure 8: Global fitting of the kinetic scheme to the measured binding and unwinding data. A-G.** Data is shown in circles and results of the model in full lines. The light-blue lines show the results of 100 bootstrap Monte Carlo runs (see Methods). **A.** Equilibrium binding of mantADP at normal conditions (75 mM NaCl) **B.** Transient kinetics of mantADP binding at normal conditions. **C.** Equilibrium binding of mantADP in the presence of 2 mM adenosine. **D.** Binding kinetics of mant-ADP in the presence of 2 mM adenosine. **E.** Unwinding rates in the presence of 2 mM adenosine. **F.** Equilibrium binding of mantADP at varying salt concentrations (75, 200, 300 mM NaCl in blue green and purple, respectively). **G.** Unwinding rates as a function of salt for low (lower) and high ATP (higher).

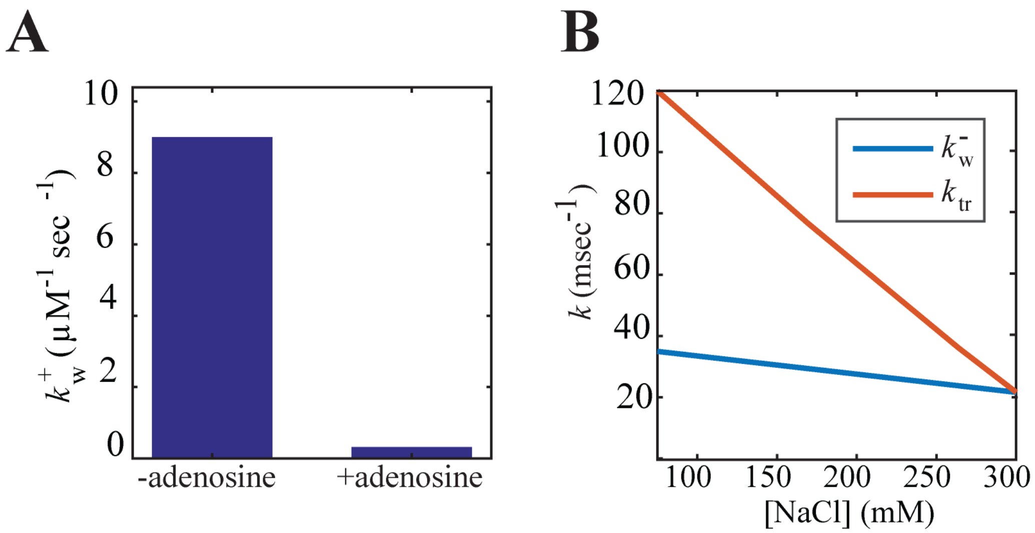


**Supplementary Figure 9: Model parameters as a function of Adenosine and salt. A.** $k_w^+$ in the presence and the absence of 2 mM Adenosine. **B.** $k_w^-, k_{tr}^+$ as a function of [NaCl].

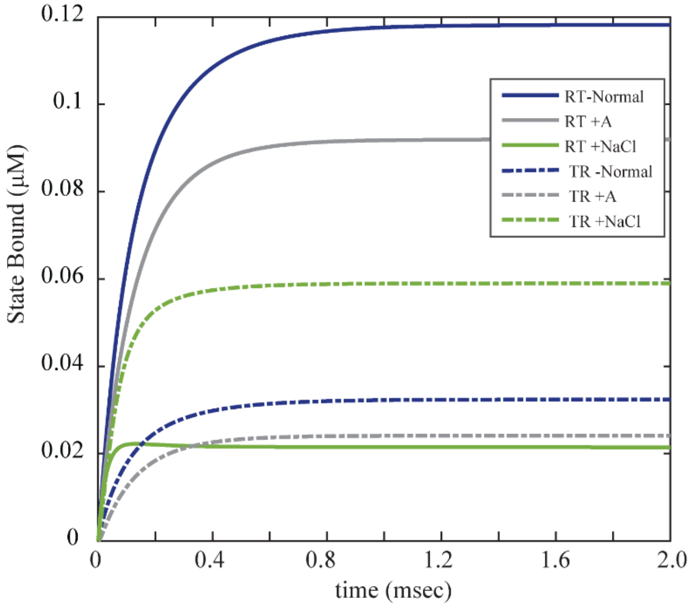


**Supplementary Figure 10: Modeled rapid mixing shows the effect of adenosine and salt on the rates and occupancies of the catalytic sites.** Rapid mixing modeling of 40 µM ATP with 1 µM RecBCD in different buffer conditions at 25 ºC (blue, 75 mM NaCl, gray 75 mM NaCl in the presence of 2 mM adenosine, and green for 300 mM NaCl).

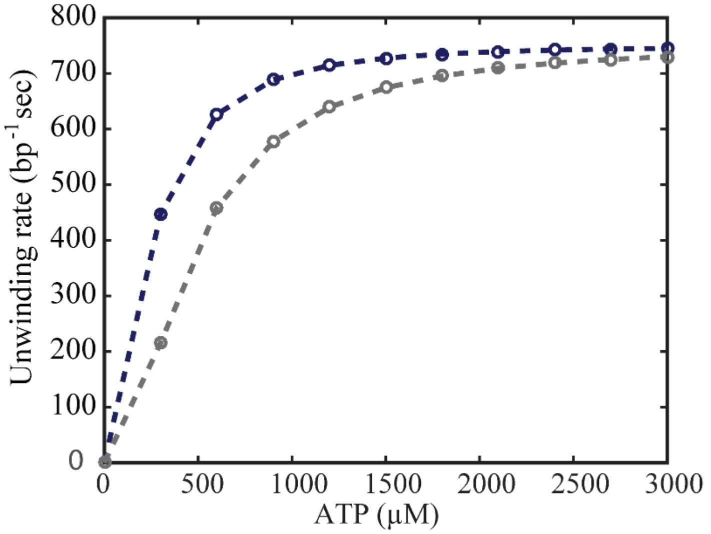


**Supplementary Figure 11: ATP-dependent unwinding rates indicate a hyperbolic-like pattern.** Simulations of ATP-dependent unwinding rate by RecBCD, in the absence (blue) and in the presence (grey) of 2 mM adenosine, indicate hyperbolic-like curves that resemble Michaelis-Menten behavior.

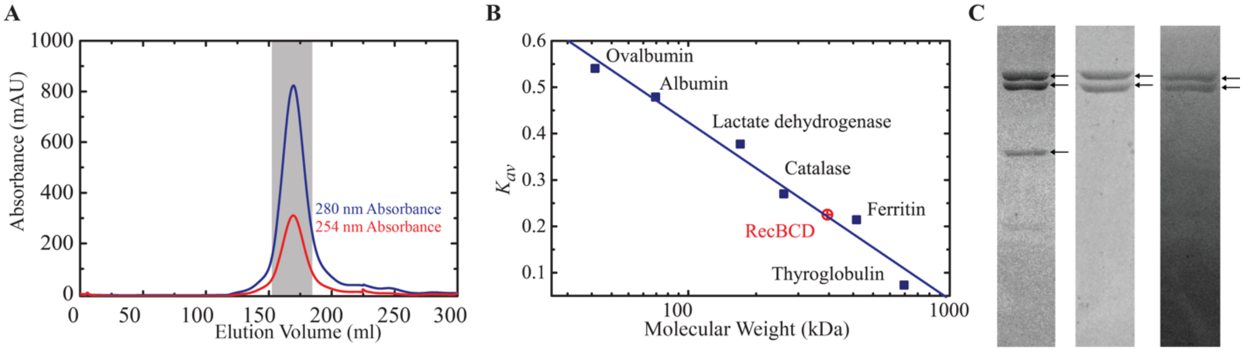


**Supplementary Figure 12: Purification of RecBCD oligomeric states A.** Size Exclusion Chromatography (SEC) analysis of RecBCD purification showing a major mono disperse peak. The major peak, which correspond to RecBCD was collected during fractionation (highlighted in grey) **B.** RecBCD predicted molecular weight according to the known standards $K_{av}$ values is calculated to be ~340 kDa. MW standards are: Ovalbumin (43 kDa), Albumin (66 kDa), Lactate dehydrogenase (140 kDa), Catalase (232 kDa), Ferritin (440 kDa), Thyroglobulin (669 kDa) **C.** Coomassie stained 10% SDS-polyacrylamide gel of RecBCD **(**left**),** RecBC (center) and $RecB^{K29Q}D$ (right) purifications. RecC, RecB and RecD are indicated with arrows.

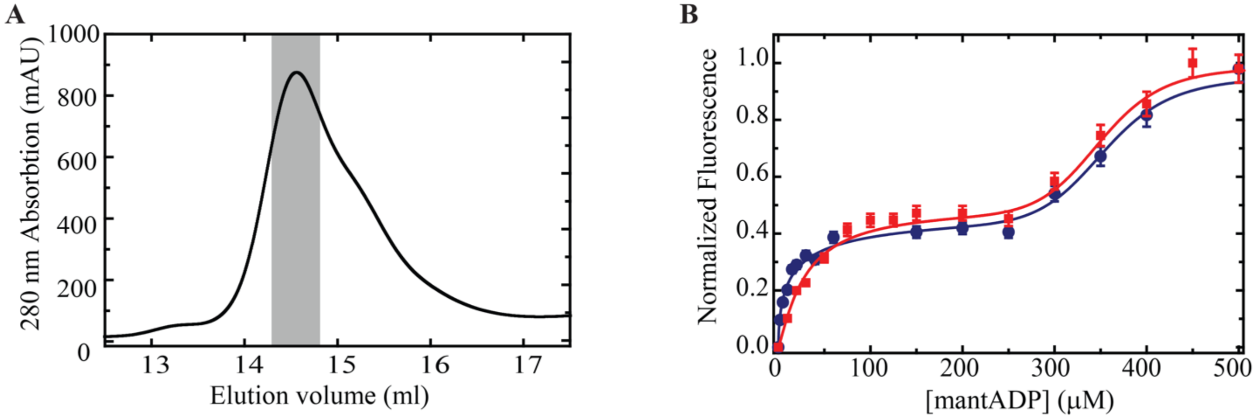


**Supplementary Figure 13: MonoQ-purified RecBCD exhibits a biphasic binding pattern.** **A.** RecBCD was purified following the same protocol as described in the 'Online Materials', immediately subjected to an additional MonoQ purification step and eluted with a salt gradient. The shaded area shows the fraction collected. **B.** The monoQ-purified protein was tested for nucleotide binding and compared with the titration in figure 1B, showing nearly identical biphasic binding.

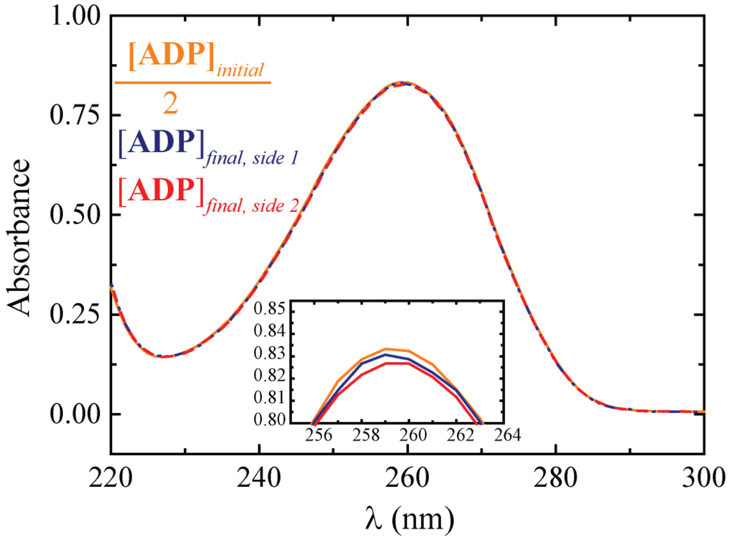


**Supplementary Figure 14: Calibration of the equilibrium dialysis experiments indicates minimal loss across the membrane.** Absorbance spectra of ADP: 20x diluted, before (yellow) and 10x diluted, after (dashed blue, side 1 and dashed red, side 2) equilibrium dialysis. The initial half absorbance at 259 nm was determined as 0.8332 (corresponding to 0.5411 M of ADP), the post-dialysis absorbance was determined as 0.8307 (0.5394 M of ADP) on the left side and 0.8267 (0.5368 M ADP) on the right side, indicating a total loss of 0.006 M of ADP and a 0.54% error. **Inset:** Zoom-in to the peaks at 259 nm.

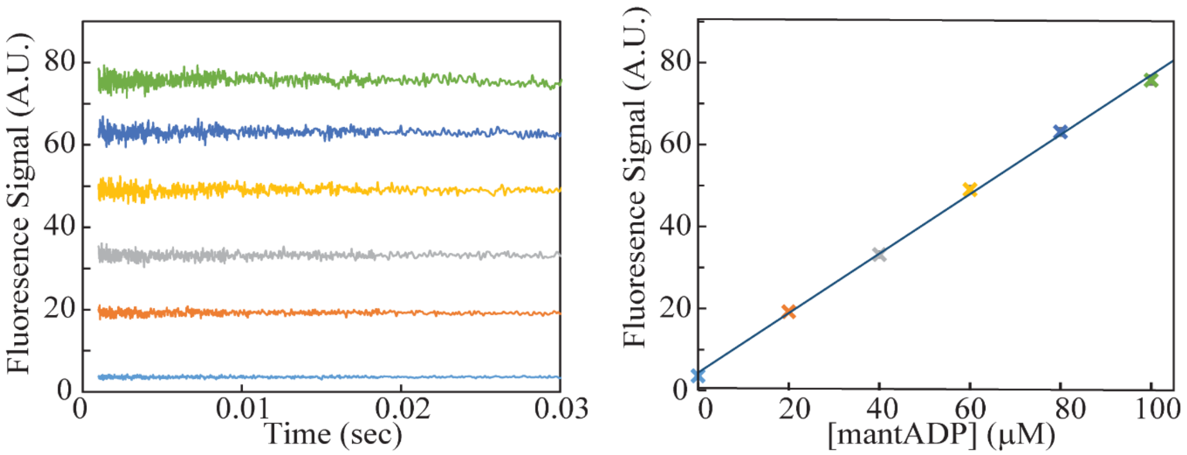


**Supplementary Figure 15: Fluorescence background quantification for binding kinetics experiments. A.** Time courses of fluorescence upon rapid mixing of buffer with mantADP (mantADP 0-100 μM, lower to upper, respectively). **B.** The time-averaged fluorescence signal displays a linear dependence on mantADP concentrations.

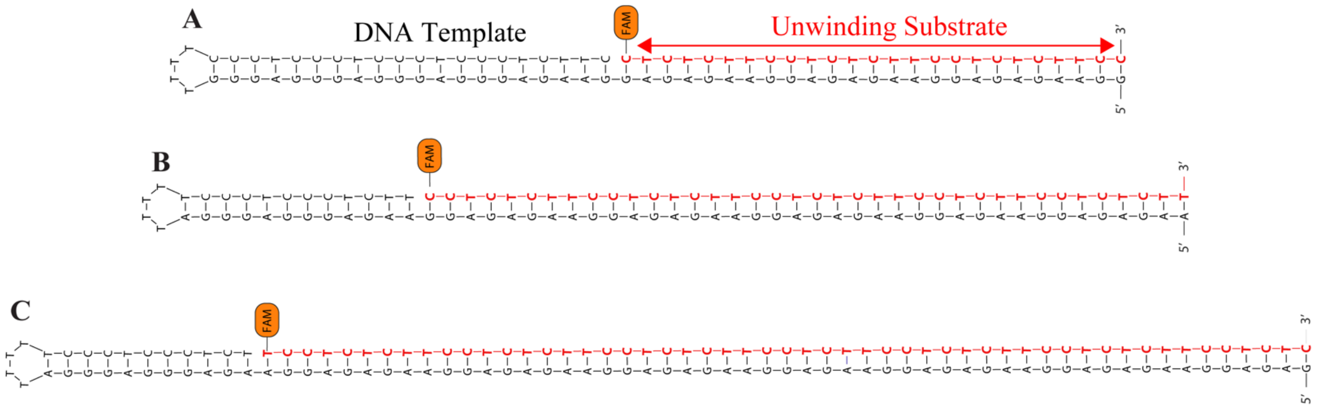


**Supplementary Figure 16**: **Schematic representation of DNA substrates used for the ensemble unwinding experiments.** 24bp (top), 38bp (middle), 52bp (bottom).

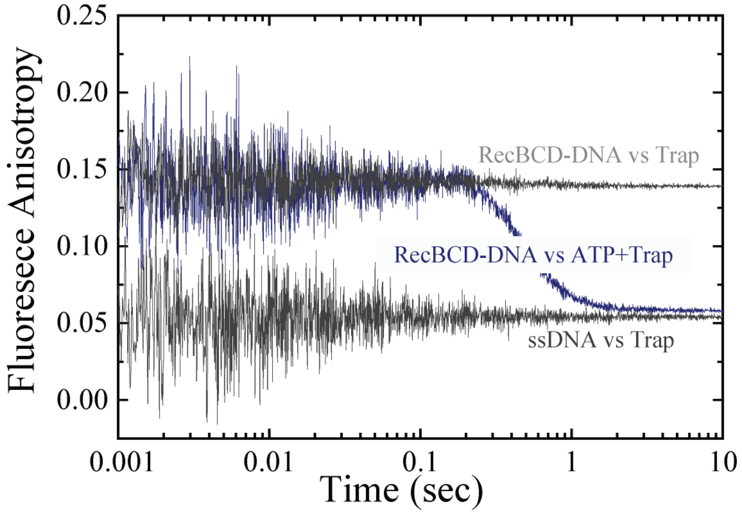


**Supplementary Figure 17: Fluorescence Anisotropy time courses reflect the physical state of RecBCD·DNA interaction.** Anisotropy time courses of rapidly mixing RecBCD·hpDNA (38bp) vs. buffer containing traps (light grey), ssDNA vs. a buffer containing traps (dark grey), and RecBCD·hpDNA vs. ATP. In the presence of ATP, the anisotropy signal drops from the value of RecBCD-bound DNA to ssDNA free in solution with a time lag representing the time taken for unwinding.

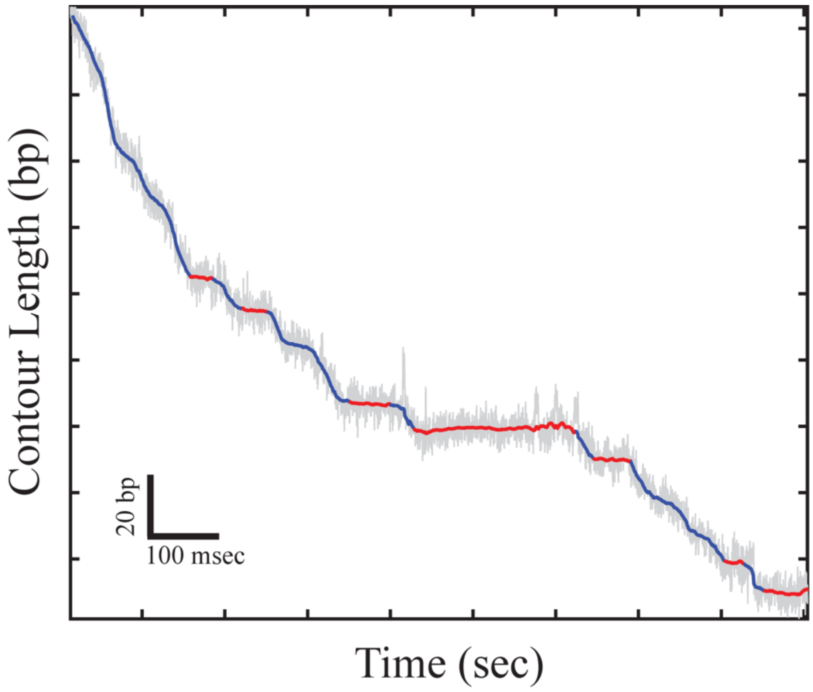


**Supplementary Figure 18: Chung-Kennedy filter and pause detection analysis of single-molecule experiments.** Representative trace (ATP 2 mM, AMPpNp 300 μM). Raw data is shown in gray. Data filtered with a Chug-Kennedy filter (Methods) is represented in blue and red; where blue represents translocation phases and red represents pauses detected by our algorithm.

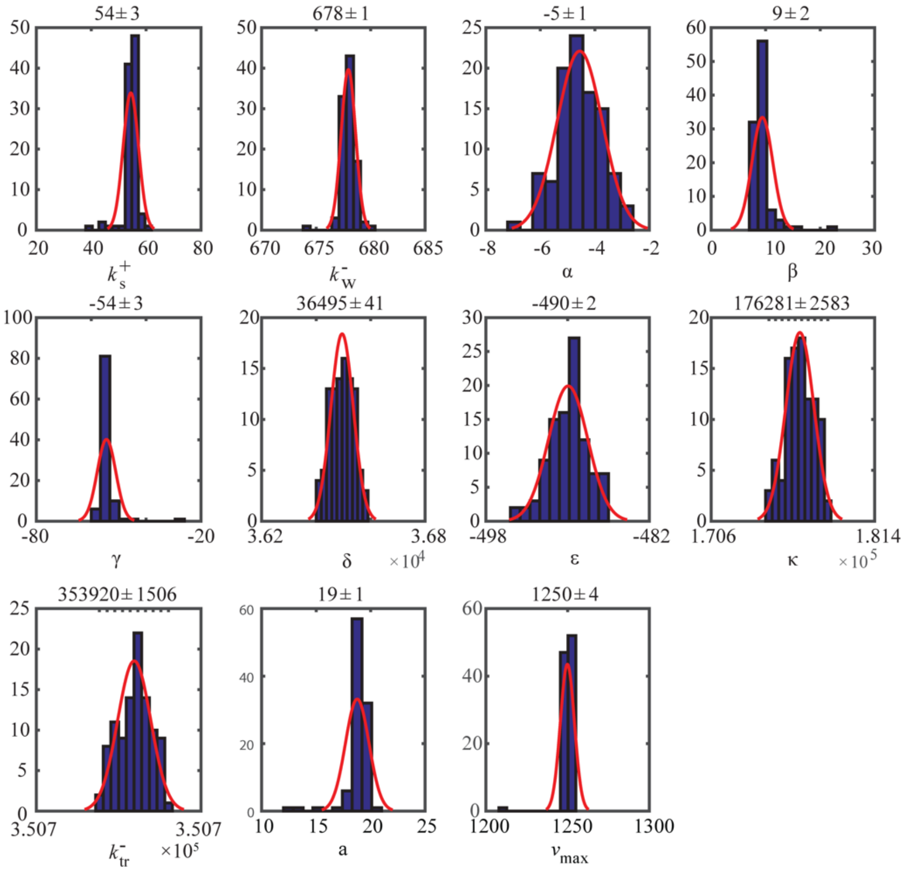


**Supplementary Figure 19: Model robustness analysis.** Distribution of the fitted parameters calculated by a Monte Carlo bootstrap test from 100 simulated datasets. Red curves through the data are best fits to Gaussian distributions. The values on top of the histograms represent their mean ± width.
